# Cancer-myeloid cell invasive program in pediatric-type diffuse high-grade glioma

**DOI:** 10.64898/2026.01.23.701142

**Authors:** Cristian Ruiz-Moreno, Raphael Collot, Thijs J.M. van den Broek, Ellen J. Wehrens, Nils Bessler, Gitanjali Dharmadhikari, Brigit M. te Pas, Ignacio L. Ibarra, Dennis S. Metselaar, Mariëtte E.G. Kranendonk, Eelco W. Hoving, Jasper van der Lugt, Friso Calkoen, Anoek Zomer, Fabian Theis, Esther Hulleman, Dannis G. van Vuurden, Anne C. Rios, Hendrik G. Stunnenberg

**Affiliations:** Princess Máxima Center for Pediatric Oncology, Utrecht, The Netherlands; Department of Molecular Biology, Faculty of Science, Radboud University, Nijmegen, The Netherlands; Oncode Institute, Utrecht, The Netherlands; PamGene International B.V., Den Bosch, The Netherlands; Institute of Computational Biology, Helmholtz Center, Munich, Germany; TUM, School of Computation, Information and Technology, Technical University of Munich, Germany; TUM School of Life Sciences, Technical University of Munich, Germany; Pediatric Oncology, Amsterdam UMC, Vrije Universiteit Amsterdam, Amsterdam, The Netherlands; Department of Biology, Faculty of Science, Utrecht University, Utrecht, the Netherlands

**Author notes:** shared first authorship (Cristian Ruiz-Moreno, Raphael Collot). shared senior authorship (Anne C. Rios, Hendrik G. Stunnenberg). Lead contact: Hendrik G. Stunnenberg.

**Keywords:** pediatric-type diffuse high-grade glioma, single-cell multi-omics, tumor microenvironment, tumor-associated myeloid cells, radial glia-like cells, invasion

## Abstract

Pediatric-type diffuse high-grade gliomas (pHGGs) are aggressive, heterogeneous brain tumors shaped by intricate cancer-microenvironment cell-cell interactions. Here, we present an integrative multimodal pHGGmap, encompassing over 800,000 cells from 136 patients profiled across transcriptomic, epigenomic, and spatial modalities. Its analysis delineated robust cancer-myeloid cell programs that structured the tumor ecosystem and identified ten distinct cancer cell states, including previously unrecognized developmental and context-responsive programs. Among these, radial glial-like (RG-like) cells exhibited dual stress-adapted and infiltrative phenotypes. Tumor-associated monocyte-derived macrophages and resident microglia engaged in four distinct immunomodulatory programs aligned with specific cancer states. Three conserved multicellular communities were maintained across treatment, including a stable, spatially and transcriptionally linked RG-like/complement-macrophage niche, indicative of cellular co-option and adaptation to support invasion. Longitudinal profiling of a metastatic diffuse midline glioma case showed that RG-like cells predominate during dissemination and remain associated with complement-enriched macrophages, whose reprogramming restores immune activation. pHGGmap establishes a landmark resource for translational discovery.

**Graphical abstract:** 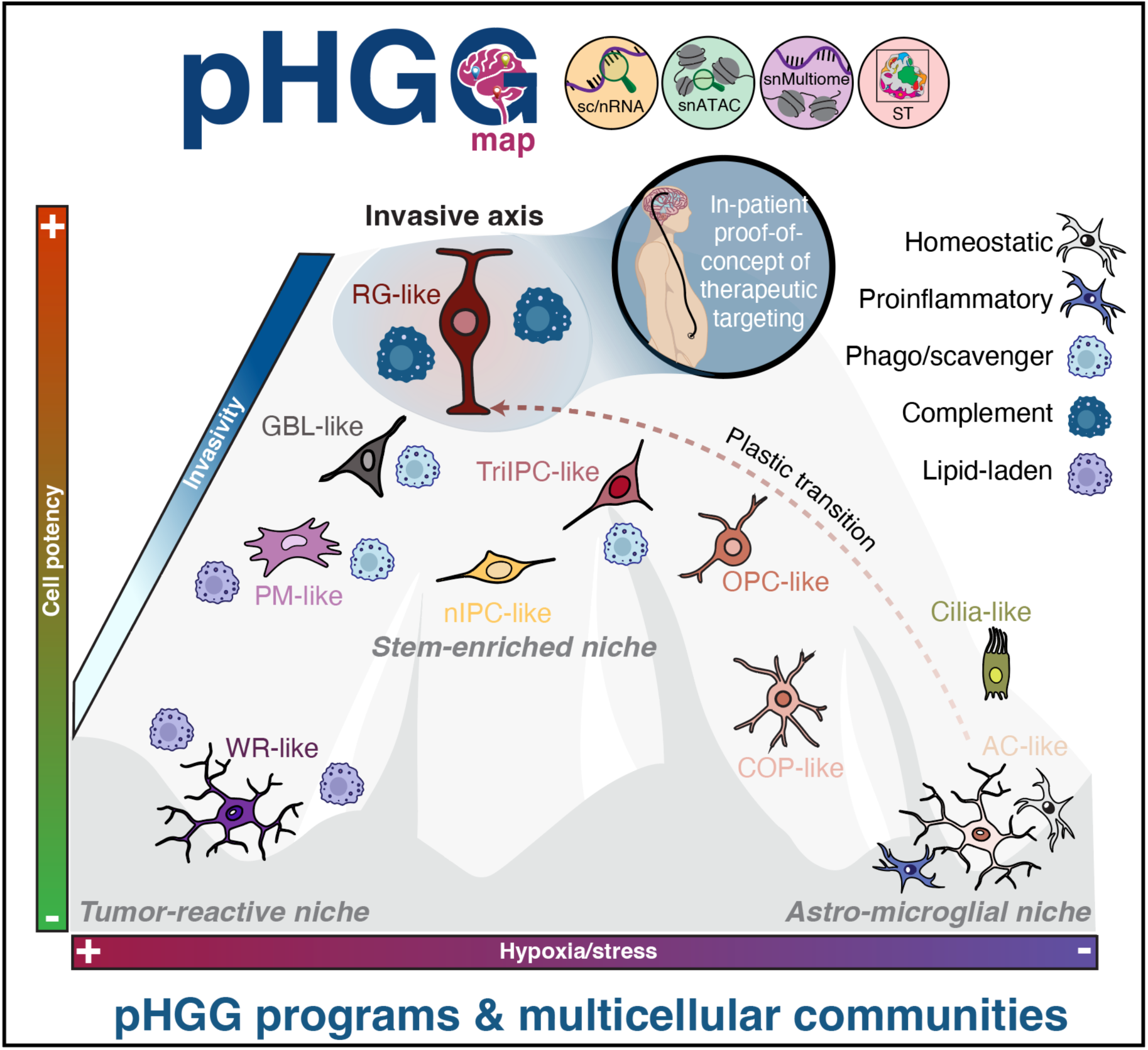

## INTRODUCTION

Pediatric-type diffuse high-grade gliomas (pHGGs) are highly aggressive and lethal pediatric brain tumors^1^. They include diffuse midline gliomas (DMGs) with H3 K27-alterations and diffuse pediatric-type high-grade gliomas H3-wild-type/IDH-wild-type (pDG-H3wt)^2^, characterized by distinct genetic alterations and preferential brain locations, featuring notable somatic and clonal heterogeneity^3^.

The dismal prognosis of pHGGs is primarily driven by their complex anatomical locations and infiltrative growth patterns, which severely limit the feasibility of surgical resection. Additionally, radio/chemotherapy, the current standard of care, offers only a marginal survival benefit, and other therapeutic strategies have consistently failed to improve outcomes^4^, although recent progress in CAR T cell therapies and oncolytic viruses offers some promise in early clinical trials^5–9^. Therapeutic failures in brain tumors are increasingly linked to their complex heterogeneity, shaped by both neoplastic and non-neoplastic cells within the tumor microenvironment (TME) and, in the case of pHGGs, driven by their early developmental origin^10,11^. The interplay between developmental programs and microenvironmental cues shapes a continuum of cellular states along axes of developmental progression and TME reactivity^12^. To develop more effective treatments and enhance tumor susceptibility to promising therapies, it is crucial to delineate cancer cell states in relation to their developmental counterparts. Malignant states may hijack key features of brain development, such as lineage plasticity^12^, migratory behavior, and immune privilege^13^, that might contribute to tumor aggressiveness. Likewise, understanding how distinct cancer cell states are organized within the microenvironment, particularly in relation to the diverse states of myeloid cells^14^ is equally important, as this may reveal conserved immune-cancer communities amenable to therapeutic targeting.

Recent studies using single-cell and spatial transcriptomic profiling have highlighted the pHGG cancer cell heterogeneity, which has predominantly been described as reflecting oligodendroglial and astrocytic signatures^15–17^, rather than the neuronal programs more commonly observed in adult tumors^12^. Furthermore, while a ‘mesenchymal (MES)-like’ signature has been identified in pediatric tumors^16,18^, additional context-specific reactive programs, primarily linked to hypoxia, have also been reported^12,17,19^. However, these reactive programs appear at lower abundance in pediatric tumors and remain less thoroughly characterized compared to those in adult counterparts^12,20^. Progress in resolving malignant phenotypes has been constrained by several factors, including reliance on murine atlases for interpreting cancer programs^21^, limited cohort sizes (both patients and profiled cells), and compartment-specific enrichment strategies that might bias the relative contributions of malignant and immune populations within individual tumors.

Here, we employed a comprehensive approach to interrogate the pHGG TME by integrating multi-omics analyses of both newly profiled and publicly available datasets, and validating the findings in external cohorts, analyzing more than 800,000 cells across 136 patients. By leveraging recently published human developmental brain atlases^22–24^, we were able to chart in detail developmental-mimicking hierarchies of cancer cells, including previously undescribed phenotypes. Our findings further clarified cellular states linked to TME factors, such as hypoxia and stress. Furthermore, we thoroughly outlined the distinct immunomodulatory programs governing tumor-associated myeloid cells (TAMCs). At both single-cell and spatial resolutions, we identified consistent multicellular communities/niches within the TME. Among the three most robust cellular communities, we identified a distinct stem-like niche enriched for radial glia-like (RG-like) cells and complement-associated TAMCs. Importantly, this niche exhibited the highest invasive potential and was validated in a patient presenting with abdominal metastasis. Our findings highlight the clinical relevance of our curated resource (pHGGmap) for translational discovery.

## RESULTS

### Multimodal Profiling of Pediatric-type Diffuse High-Grade Gliomas

To comprehensively decode the pHGG cellular landscape, we performed multimodal profiling integrating data from single-cell/nucleus RNA sequencing (sc/nRNA-seq), single-nucleus ATAC-seq (snATAC-seq), single-nucleus Multiome (snMultiome), and spatially resolved transcriptomics (ST) (**Graphical abstract**). Our discovery cohort integrates transcriptomics and epigenomic information from 73 patients, including 37 newly profiled patients and 36 from published studies^16,17^ (**Figures 1A and 1B; Table S1**). Following quality control, background noise removal, and doublet filtering, we obtained 466,778 cells (342,184 transcriptome and 124,594 ATAC) (**Figures 1A, 1C, S1A and S1B**). In addition, we included a validation cohort of 63 cases^15,18,20,25,26^, comprising 350,612 cells, to independently corroborate and extend our primary findings (**Figure 1A; Table S2**).

**Figure 1.**
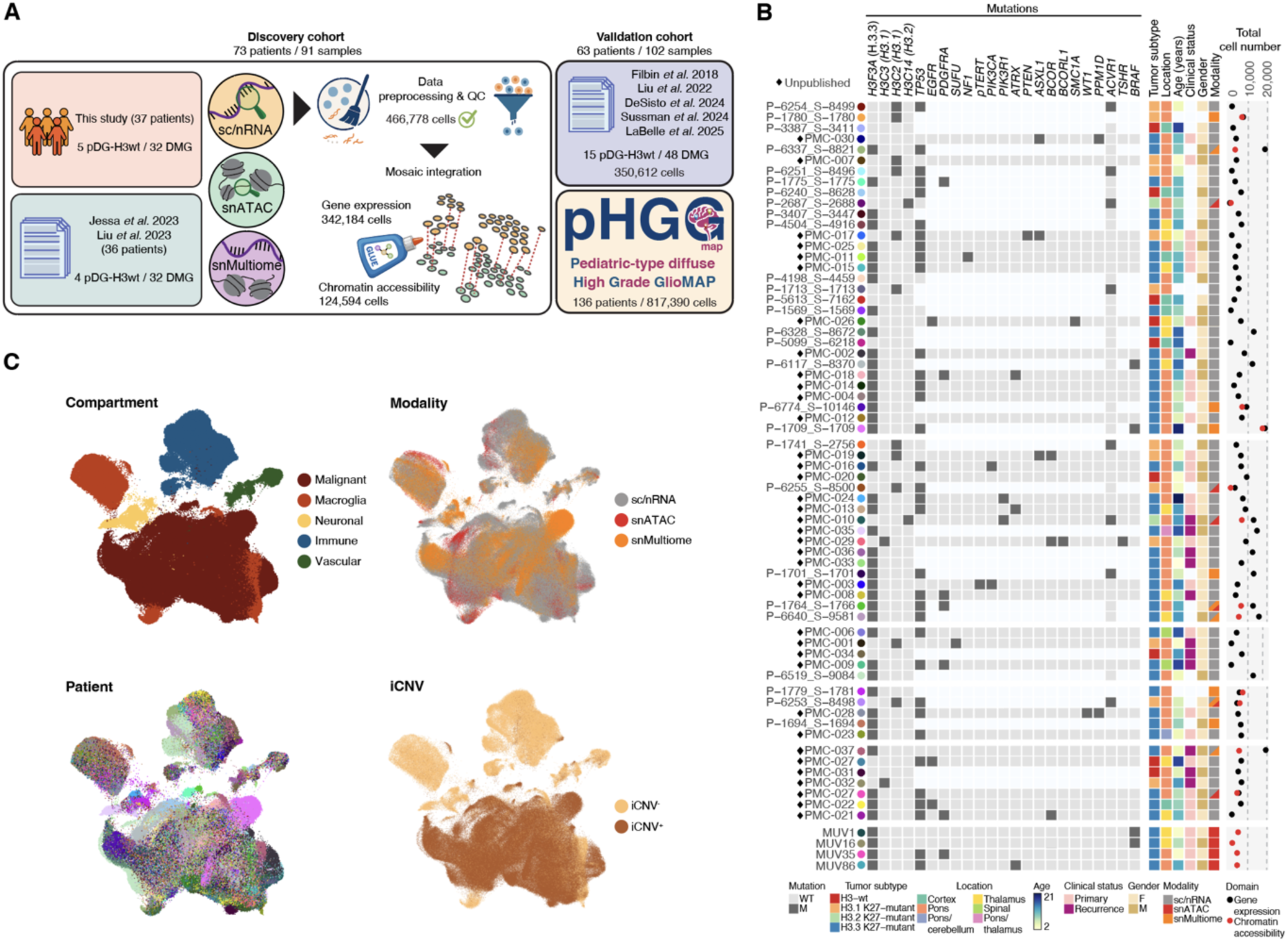
Multimodal map of pHGG - pHGGmap. (A) Schematic of the study design. (B) Patient’s clinical and molecular features (pHGGmap - discovery cohort). Patients are ordered according to cellular composition, as in **S1F**. (C) UMAP representation of the discovery cohort colored by cell compartment, modality, patient (color code in **1B**), and iCNV. See also **Figure S1**.

Because multimodal datasets often contain substantial batch effects, we evaluated three data integration methods using the single-cell integration benchmark pipeline^27^ (**Figure S1C**). Quantitative evaluation of low-dimensional representations revealed that GLUE^28^ offered the best balance between batch correction and biological conservation (**Figures 1C and S1C**). Malignant cells were detected using a high-resolution, haplotype-aware inferred copy number variation (iCNV) method^29^ (**Figures 1C and S1D**). The integrated pHGGmap (discovery cohort) comprised five main compartments, including malignant (iCNV⁺) cells and the surrounding microenvironment (iCNV⁻), composed of macroglia, neuronal, immune, and vascular cells (**Figures 1C and S1E**). Differences in the proportional representations of cell subtypes, such as malignant-, immune-, and neuro/glial-enriched (**Figure S1F**), revealed distinct patterns across patients, underscoring the need for large cohorts to prevent over-generalization of the tumor cellular landscape in pHGG, and enabling the detection of the most promising and biologically relevant patterns.

### Defining cancer meta-programs

We next sought to resolve the spectrum of cancer cell phenotypes by applying non-negative matrix factorization (NMF) to identify gene programs that vary among cells in each patient, as previously described^30^ (**Figure 2A)**. This analysis revealed 552 robust NMF programs. Hierarchical clustering identified recurrent expression patterns, from which we derived 13 meta-programs (MPs) (**Figure 2B; Table S3**). The MPs encompass diverse cellular processes, including core biological functions, such as cell cycle (MP1), hypoxia (MP6), and stress responses (MP10), previously identified across pan-cancer single-cell transcriptomic datasets^30–32^. By comparing our results to previously described glioma gene signatures^17,20,32–36^ (**Table S4**), we identified seven neuroglial-like programs and three programs that segregate with pan-cancer epithelial-to-mesenchymal transition (EMT)^30^ and MES glioma signatures^20,31,33,34^ (**Figure S2A**), establishing a framework for interpreting intra- and intertumoral cancer states across pHGG patients.

**Figure 2.**
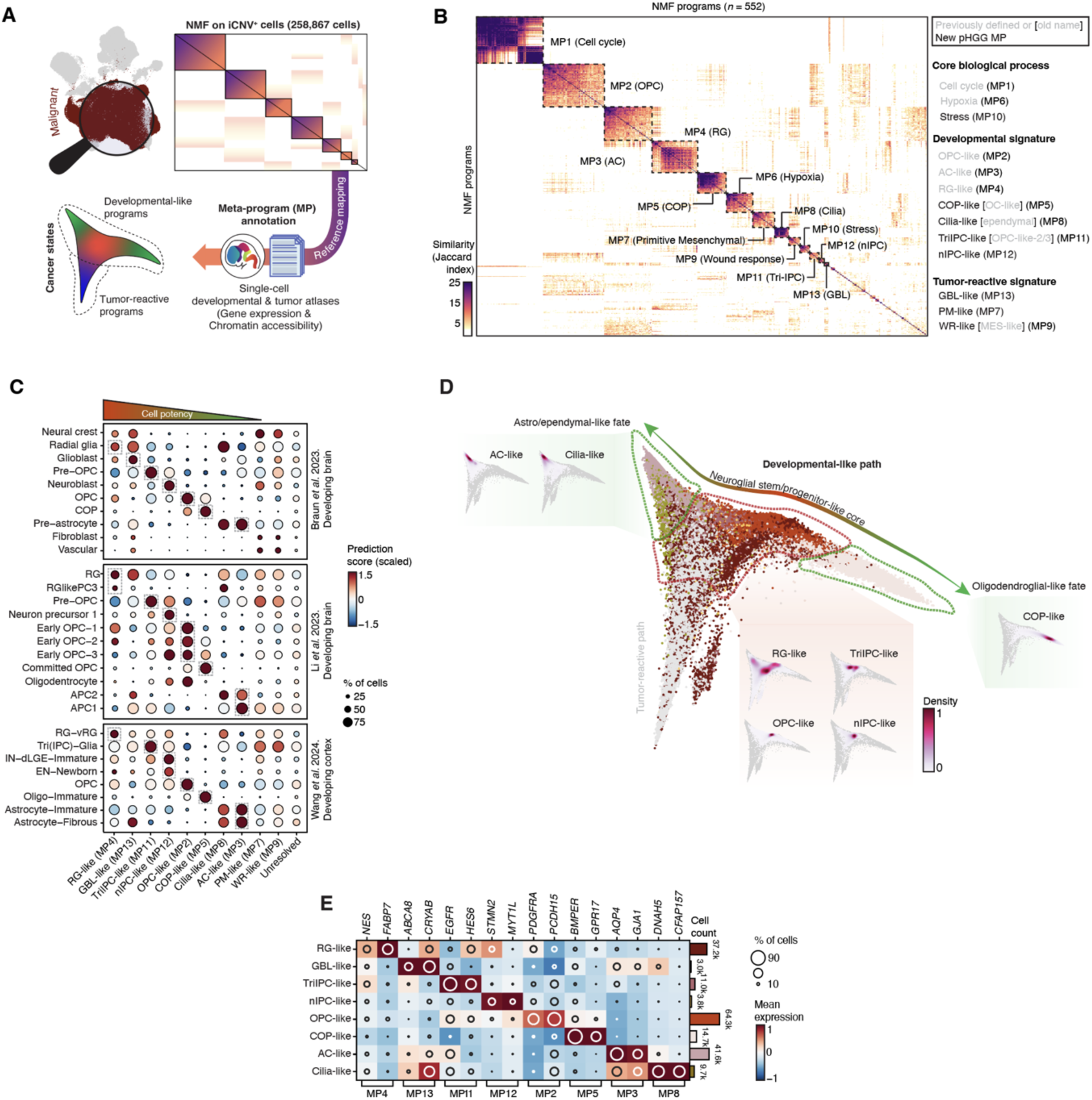
Characterization of developmental cancer cell states. (A) Schematic of the analytical pipeline. (B) Heatmap shows the Jaccard similarity indices for comparisons among robust NMF programs based on their top 50 genes. (C) Dot plot displays prediction scores of pHGG cancer cells (per MP) after mapping onto reference atlases^22–24^. Original annotation per reference was kept. (D) Diffusion map (DC1-DC3) of malignant cells colored according to the higher matching developmental MP and density. Grey-colored cells did not have a brain developmental cell match. (E) Expression of selected markers per developmental MP. See also **Figure S2**.

### A spectrum of human neurodevelopmental-like cancer programs

Building on these MPs, we leveraged three detailed atlases of human brain development^22–24^ to explore neuroglial lineage-specific trajectories in greater detail (**Figure 2C**). Ordering the cells’ transcription profiles by their similarity in a diffusion map revealed a tumor pseudo-hierarchy spectrum, uncovering transitions between cancer cell identities that closely mirror developmental trajectories **(Figures 2C and 2D).** Consistent with previous findings^15–17^, we identified gene programs that align with differentiated glial cells and progenitor-like cancer states **(Figures 2C and 2D).** Among the more differentiated states, MP5 exhibited signature overlap with the oligodendrocyte-like phenotype^15,16^ **(Figure S2A)**. However, reference mapping to human developing brain cell atlases showed that MP5-expressing cells are better classified as committed oligodendrocyte precursors (COP-like), characterized by high expression of *BMPER* and *GPR17*, which are absent in mature oligodendrocytes^22^ **(Figure 2E and S2B)**. Our observations were further supported by *in silico* mixing of developing and adult brain references^37^ (**Figure S2C**). MP3 exhibited a continuum of features spanning both immature and mature astrocytes (AC) (**Figure 2C**), forming a gradient rather than distinct precursor and differentiated states, which led us to retain the AC-like nomenclature. In addition, we found that MP8 was enriched in the CFAP gene family, which encodes cilia- and flagella-associated proteins (**Figure 2E**). This program overlaps with the ependymal-like cells identified in H3.1 DMG^17^ **(Figure S2A)**. However, we demonstrate that this program is not limited to H3.1 but is also present in H3.2 (a rare variant) and H3.3 DMG, as well as in pDG-H3wt (**Figure S2D**).

Among the most undifferentiated states, MP2 (*PDGFRA*^hi^) mirrors oligodendrocyte precursor cells (OPC-like) **(Figures 2C, 2D, and 2E)**, representing a stalled transitional proliferative state, well-recognized as a central driver of tumorigenesis, particularly for the DMG subtype^15,20^. MP12 and MP11 are closely associated with MP2 (OPC-like) (**Figure S2A**) yet present more neural precursor attributes. Specifically, MP12 enriched for genes linked to early neural fate, including *STMN2* and *MYT1L*, consistent with neural-committed intermediate precursor cells (nIPC-like) **(Figures 2C, 2D, and 2E**) and closely correlates with NPC2-like^34^ and NPC Glioma^30^, identified in adult glioma (**Figure S2A**). MP11 (*EGFR, HES6*) resembled the pre-OPC phenotype **(Figure 2C and 2E)**. This aligns with previously identified populations of *EGFR*^hi^ cells, which give rise to OPCs or mature glial cells and have been designated as OPC-like2**/**3 in DMG^20^. However, recent work by Wang *et al.*^23^ demonstrated that *EGFR*^hi^ cells can locally differentiate not only into OPC and AC but also into neuronal precursor cells (e.g., nIPC), redefining them as tripotential intermediate progenitors (Tri-IPC), revealing a multipotent state that challenges traditional neuronal and glial lineages. The discovery of nIPC-like state in pHGG led us to reclassify (pre)OPC-like 2/3 cells^20^ as potentially being in a Tri-IPC-like malignant state, suggesting that these subsets may retain broader lineage flexibility. At the most primitive end of the developmental spectrum, MP4 (*NES, FABP7*) exhibited substantial similarity to radial glial (RG) cells, aligning with predictions from multiple developmental datasets **(Figures 2C and 2E)**. While an RG-like cancer state was previously proposed^19^, our approach offers the first detailed characterization of this population within the tumor hierarchy. Our MPs were further confirmed in the validation cohort^15,18,20,25^ **(Figures 2F and S2E)**. Collectively, our analysis revealed an expanded neurodevelopmental-mirroring continuum in pHGG, identifying four stem/progenitor-like states, including RG- and nIPC-like cells, with refined branching trajectories toward astrocytic/ependymal and oligodendroglial-like lineages. In addition, we proposed a refined characterization of tumor states, including the reclassification of mature oligodendrocyte-like^15,16^ as COP-like and the OPC-like2/3^16^ populations as TriIPC-like states, deepening our understanding of tumor cell lineage diversification in pHGG.

### Tumor-reactive cancer states

A continuum of cell states, ranging from developmental to environmentally induced MES-like phenotypes, has been described in glioma^12^, with one MES-like program previously identified in pHGG^18,20^. Consistent with this, we identified three programs: MP13, MP7, and MP9, enriched for hypoxia and stress signatures, indicating they might be shaped and react to TME conditions (hereafter tumor-reactive) **(Figures S2A and S3A; Table S3**). MP13, a signature not previously reported in pHGG, resembled glioblasts (GBL-like) with high *ABCA8* and *CRYAB* expression^22^ (**Figures 2C and 2E**) and also matched the fibrous astrocyte signature from Wang *et al.*^23^, suggesting a transient, environment-induced state along the AC-like developmental trajectory. In contrast, MP7 and MP9 lacked transcriptomic enrichment to any neuroglial cell type during early brain maturation (**Figure 2C**). MP9 showed similarity with the MES-like state described in adult GB and DMG^18,20^ (**Figure S2A**). However, Gene Ontology (GO) analysis revealed enrichment of biological processes involved in the inflammatory response, wound healing, and apoptosis, with a lack of terms associated with MES-like states (e.g., EMT)^38^ (**Figure S3B**). In addition, SCENIC+ analysis^39^ shows a set of transcription factors (TFs) that differs from those related to neurodevelopment and leans towards immune-like (BATF3)^40^ or damage responses (BCL6)^38^ (**Figure S3C**). This aligns with recent work by Mossi Albiach *et al.*^41^, challenging the definition of the MES-like program in adult GB and redefining it as a wound response phenotype (WR-like). The identification of the WR-like state as an inducible functional state that responds to environmental signals, such as hypoxia or tissue stress^41^ aligns with a higher level of epigenetic plasticity of this state based on our analysis employing a classifier^42^ based on chromatin accessibility near genes rather than gene expression alone (**Figure S3D**). On the other hand, MP7, which also exhibits a high ‘MES-like’ overlap (**Figure S2A**), lacks classical reported genes (e.g., *CD44*) found in the WR-like gene signature (**Figure 3B**). Instead, MP7 is enriched for genes associated with a more primitive state, such as *HMGA2, COL19A1,* and *ITGA3* (**Figure 3C**). GO analysis suggests that MP7 is linked to processes involving other germ layers during embryonic development and *bona fide* EMT hallmarks (**Figures 3C and S3E)**. Consequently, we refer to this state as primitive-MES like (PM-like). Because of this primitive-like identity, we explored the stem-like potential of PM-like cells by employing a deep learning framework for characterizing potency and differentiation states (CytoTRACE2)^43^. While, as expected, RG-like cells exhibited the highest stemness potential, we found that PM-like cells ranked second, surpassing other neuroglial progenitor-like cancer cell states, including TriIPC- and OPC-like cells. (**Figure 3D**). These findings point to a scenario in which some cancer cells might either retain a primitive, developmentally immature non-neuroglial phenotype or transdifferentiate into such a state in response to environmental cues. We observed increased enrichment of genome accessibility patterns characteristic of endoderm (e.g., esophageal carcinoma [ESCA]) or mesoderm-derived (e.g., mesothelioma [MESO]) tumor types (**Figure 3E**), further supporting the divergent nature of this program in relation to the WR-like. Furthermore, analysis of the enhancer-driven gene regulatory networks from PM-like cancer cells revealed TFs that are critical for early development, stem cell renewal, maintenance of pluripotency, and early lineage commitment, such as FOSL1^44,45^, HMGA2^46^, and E2F7^47^ (**Figure S3F and S3G**). Predicted target genes regulated by these TFs, such as *SALL4*^48^, *IGFBP2*^49^, and *ITGA5*^50^, are known to play key roles in stem cell maintenance, early lineage specification, and pluripotency, particularly in mesoderm and early tissue development. Our analyses allowed us to uncover the distinct nature of the PM-like cancer state. Moreover, examining the expression of its key defining markers revealed its absence in high-grade gliomas in adults (**Figure 3F**). Altogether, we defined three reactive cell states in pHGG, characterized by a strong connection with environmental factors - hypoxia and stress - revealing novel cancer phenotypes, including PM-like, with high stem-like potential, unique to this tumor type.

**Figure 3.**
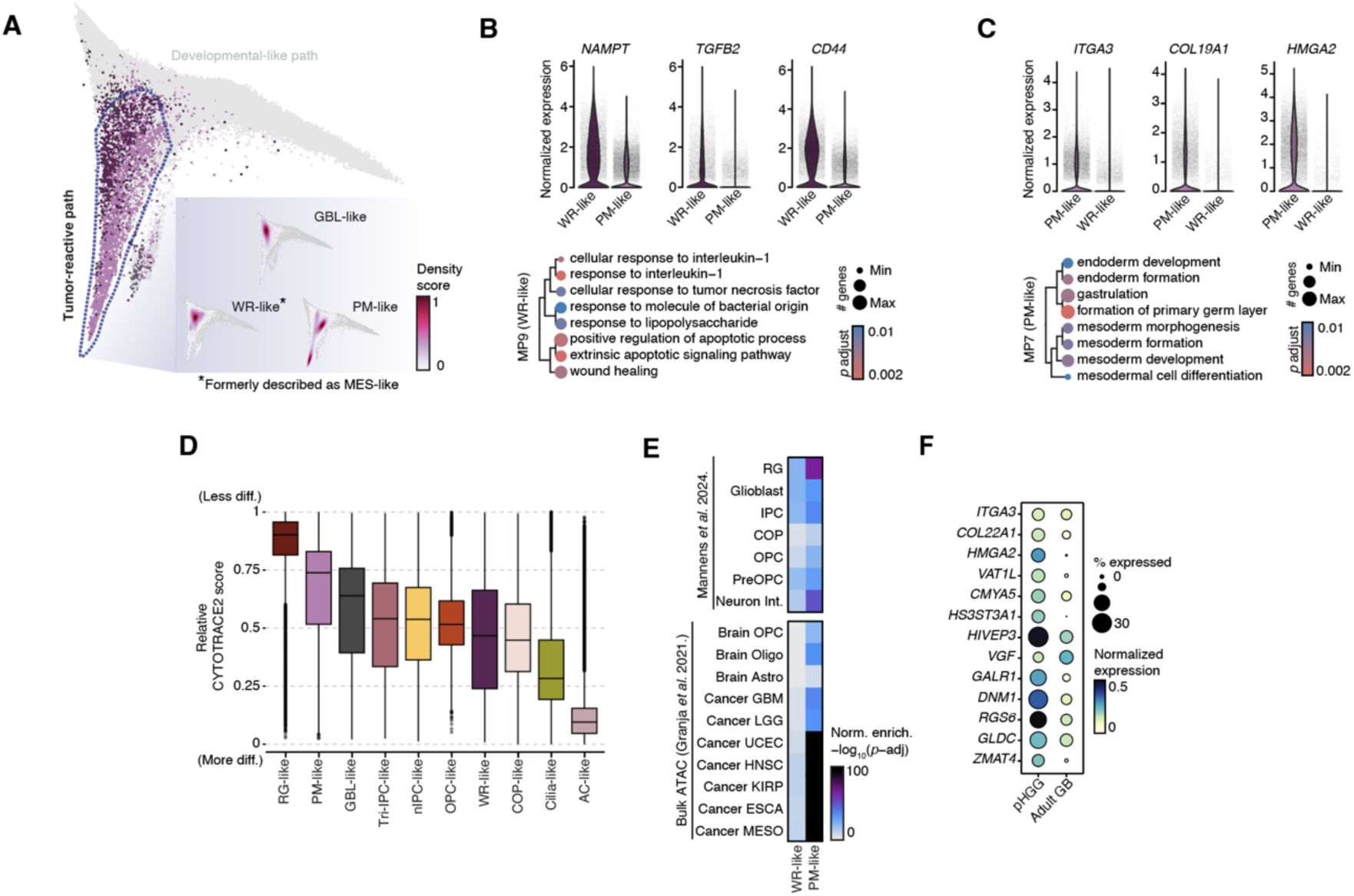
Tumor-reactive programs. (A) Diffusion map (DC1-DC3) of malignant cells colored according to the higher matching tumor-reactive MP and density. Grey-colored cells correspond to the brain developmental-like cancer MPs. (B) Gene expression of genes enriched in the WR-like cells. Bottom, a dot plot of the top significant pathways per MP from the GO:BP database. (C) As in (B) but in the PM-like cells. (D) Box plot of normalized potency scores determined by CytoTRACE2^43^ from 0 (more differentiated) to 1 (less differentiated). (E) Heatmap showing normalized enrichment scores of single-cell (top)^95^ and bulk ATAC-seq (bottom)^96^ references across tumor-reactive states. (F) Expression of the top PM-like markers in pHGG and adult GB. Also see **Figure S3**.

### Monocyte-derived macrophages and resident microglia engage distinct immunomodulatory programs

As previously reported, immune cells, particularly tumor-associated myeloid cells (TAMCs), represent the most abundant non-malignant component in the pHGG TME (**Figure S1F)**, displaying diverse phenotypes^26,51,52^, likely shaped by the brain’s developmental context, tumor-intrinsic programs, and local inflammatory signals. Applying the NMF approach to myeloid cells in the discovery cohort, we identified 358 robust programs across patients, defining after hierarchical clustering, 13 consistent MPs (**Figure 4A and B; Table S5**). We compared these gene modules to published signatures from immune-focused studies in cancer^14,30,53–56^ to delineate cell identity, core biological processes, and functional activities (**Figures 4A and S4A; Table S6**). Among these programs, only two displayed distinct cell identity signatures. MP4 featured classical monocyte markers (*LYZ, FCN1*) along with genes associated with infiltration and early activation (*VCAN, IL1R2*) (**Figure 4C**). In contrast, MP10 was enriched for canonical markers of resident microglia (MG), including *P2RY12* and *CX3CR1* (**Figure 4C**). Consistent with our previous findings^52^, these cells (20,6% of TAMCs) retained core homeostatic microglial gene expression and lacked transcriptional features associated with disease-associated MG^57^, as indicated by the absence of metabolic gene upregulation (*IGF1, FABP5, FLT1*)^58^ and downregulation of homeostatic markers (*P2RY12*, *CX3CR1*)^58^ **(Figures S4B and S4C**). Homeostatic MG shows a significant association with the synaptic pruning program (MP6) **(Figure S4D)**, a key function in preserving neuronal integrity. Both MPs aligned closely with the homeostatic microglial signature described in a non-tumor brain atlas^59^ **(Figure 4D)**. This further supports the notion that most tumor-embedded MG in pHGG retain functional similarities to their healthy counterparts.

**Figure 4.**
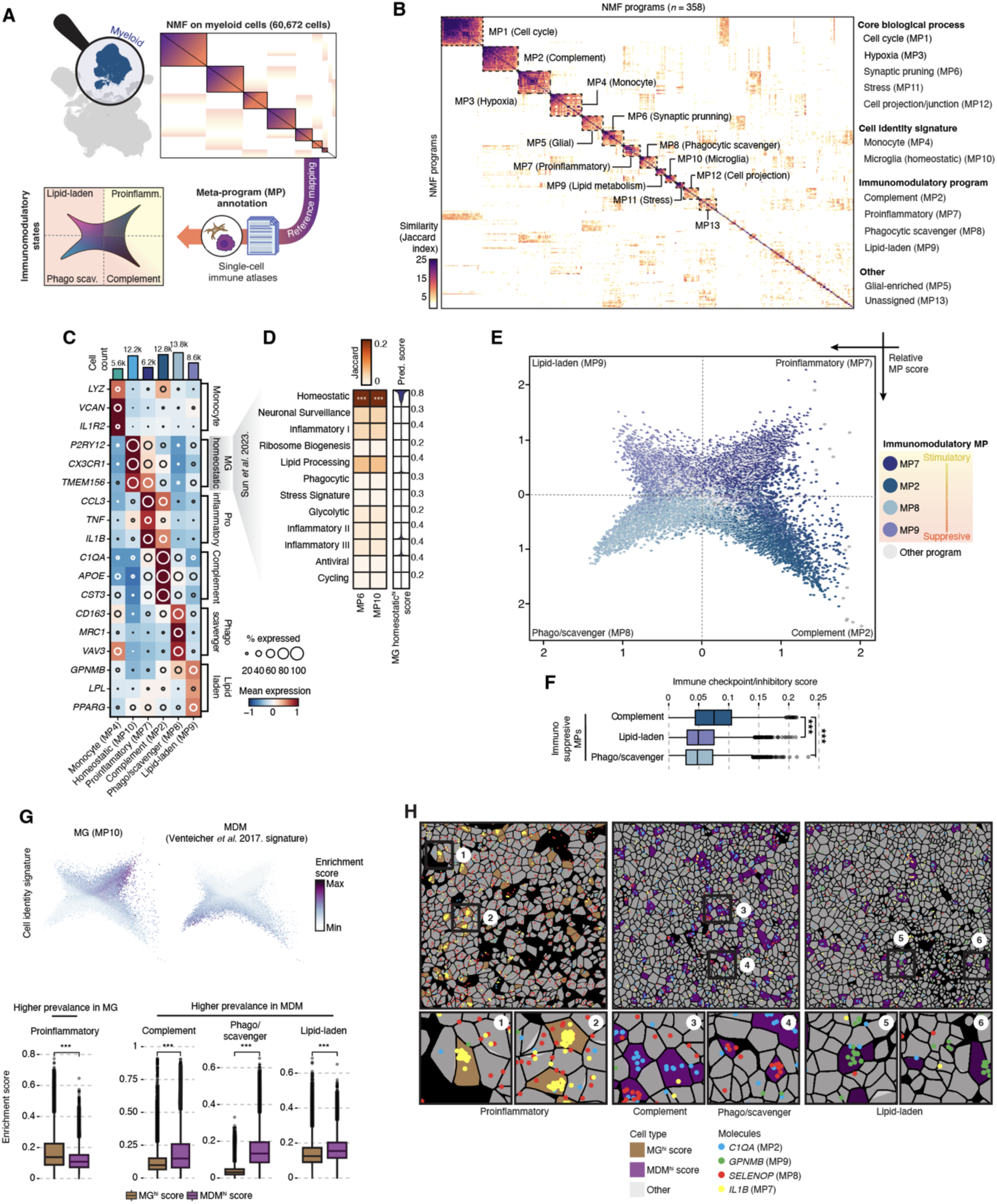
Non-, pro-, and anti-inflammatory programs in TAMCs. (A) Schematic of the analytical pipeline. (B) Heatmap shows the Jaccard similarity indices for comparisons among robust NMF programs based on their top 50 genes. (C) Expression of selected markers per MP. (D) Jaccard similarity heatmap illustrating the significant correspondence of marker genes between Sun *et al.*^59^ and MP6 and MP10. The significance of the overlap is assessed using Fisher’s exact test. \*\*\**p* < 0.001. Right, violin plot displays the prediction score when performing reference mapping using Azimuth^97^. (E) Representation of cell states. Every quadrant represents a distinct immunomodulatory TAMCs MP, and each dot, structured as a pie chart, showcases the distribution of MPs within each cell. (F) Box plot shows the score of the immune checkpoint/inhibitor genes. Statistical significance was assessed using a Wilcoxon test. \*\*\**p* < 0.001. (G) Enrichment scores of selected cell identity MPs. Below, box plots show the enrichment score of immunomodulatory MPs for MDM and MG. Statistical significance was assessed using a Wilcoxon test. \*\*\**p* < 0.001. (H) Selected fields of view (FOVs) from van den Broek *et al.* CosMx dataset^65^ displaying subcellular expression map of marker genes that define each immunomodulatory MP. TAMCs are colored by ontogeny, whereas other cells are shown in grey. See also **Figure S4**.

Although none of the meta-programs matched a ‘naïve’ monocyte-derived macrophage (MDM) ontogeny, we identified TAMCs that lacked MG markers and instead expressed canonical MDM-associated genes, including *ITGA4, TGFBI*, and *CD163*^60,61^. Notably, TAMCs with high MDM signature scores were almost exclusively enriched in hypoxia (MP3) and stress (MP11) programs **(Figure S4D),** suggesting a strong influence of the TME on their state. After identifying the main myeloid ontogenies, we examine their relationship to the four immunomodulatory programs revealed in our analysis (**Figure 4E**). We identified a single pro-inflammatory program (MP7), present in 10,5% of the TAMCs and primarily associated with MG (**Figures 4F and S4B**). MP7 was marked by inflammatory cytokines and chemokines involved in lymphocyte and monocyte recruitment (e.g., *IL1B, CCL4L2, CCL3, CCL4, TNF*), along with immediate early genes (*EGR1, EGR2*)^62^ (proinflammatory-TAMCs, **Figures 4C and S4E; Table S5**). MP8 shared common genes with previously described scavenger immunosuppressive^14^ and phagocytic activity programs^53^, including *CD163, CD163L1,* and *MRC1* (phago/scavenger-TAMCs, **Figures 4C and S4E; Table S5**). MP9, linked to lipid metabolism, included genes such as *GPNMB, LPL, PPARG,* and *SPP1*, and resembled the expression profile of recently described pro-tumorigenic lipid-laden TAMCs in adult GB^63,64^ and pHGG^65^ (lipid-laden-TAMCs, **Figures 4C and S4E; Table S5**). Lastly, MP2, a complement-enriched program (*C1QA, C1QC*), featured genes associated with an anti-inflammatory phenotype, such as *APOE, CST3,* and *FTL* (complement-TAMCs, **Figure 4C; Table S5**). It also showed increased expression of MHC-II antigen-presentation genes (*HLA-DRA/DRB1/DPA1/DPB1, CD74*) and associated biological pathways (**Figure S4E**). Our complement program correlated with recently described ‘complement immunosuppressive’ macrophages in adult glioma^14^ and several pan-tumor immune-related gene signatures^30,56^ (**Figure S4A**). Among the immunosuppressive programs, the complement-TAMCs exhibited a higher score and expression of checkpoint and inhibitory genes such as *HAVCR2* and *LGALS9*, supporting a key immunoregulatory role in the pHGG TME (**Figures 4F and S4F**). However, these TAMCs also expressed to some extent pro-inflammatory genes (e.g., *CCL3*) (**Figure 4C**), further suggesting a context-dependent state, in which cells might retain the capacity to respond to stress or damage signals within the TME while supporting tumor growth and spread. All three immunosuppressive programs were detectable across TAMCs, but their expression was skewed toward MDM cells, most notably the phago/scavenger program, which appeared almost exclusively in MDMs (**Figure 4G**). The preferential pro- and anti-inflammatory phenotype of MG and MDM, respectively, was validated in tissue using our previously ST dataset^65^ (**Figure 4H**) and further corroborated in the validation cohort^18,20,25,26^ (**Figure S4G**). Altogether, we defined distinct TAMC molecular programs that reveal unique features in pHGG compared to other high-grade gliomas^14^, with MGs and MDMs preferentially engaging in separate immunomodulatory pathways, indicating both cellular specialization and context-dependent adaptation within the TME.

### Therapy-associated remodeling of cancer and myeloid cell states

Having thoroughly characterized cancer and myeloid cell states, we proceeded to evaluate whether clinicopathological and/or technical factors shaped cellular composition in pHGG using a Poisson linear mixed model^66,67^. We found that the treatment status (treatment-naïve vs. post-treatment) was the most significant variable in explaining the variance, followed by capture modality (cells vs. nuclei) and tumor type (**Figure 5A and Technical note**). To focus on therapy-related effects, we reannotated and integrated cells from the Sussman *et al.* study^18^, which longitudinally profiled cases of pHGGs. Using label transfer with our pHGGmap as a reference, we harmonized cell identities to generate a treatment-focused pHGG dataset comprising 433,649 cells from 51 patients (104 samples, including 14 paired cases; **Figure 5B-i**). In this dataset, all analyzed cells were profiled using single-nuclei transcriptomics, overcoming the potential influence of the capture modality as a source of cell compositional differences. Because tumor type still explained significant residual variance, we analyzed changes in tumor composition upon therapy in DMG and pDG-H3wt, separately. pDG-H3wt showed no discernible differences between treatment-naïve and post-treatment samples, potentially due to the smaller number of cases compared to DMG (16 vs. 35 patients) (**Figure 5C**). DMG, by contrast, displayed pronounced shifts in both cancer and myeloid compartments. AC-like cells were more prevalent in treatment-naïve cases. Mirroring changes reported in adult GB^68,69^, WR-like cells showed an increase in post-treatment samples (**Figures 5C and S5A**). Consistent with recent research in pHGG^26^, we found that the most striking differences lay in the composition of the TAMCs (**Figure S5B**). Treatment-naïve samples mainly exhibited MG-associated homeostatic and pro-inflammatory programs. In contrast, following therapy, we observed a marked shift in the TAMC landscape, now characterized by the dominance of early infiltrating monocytes and TAMCs expressing immunosuppressive phago/scavenger and lipid-laden programs, commonly associated with MDMs (**Figures 5C, S5A and S5B**).

**Figure 5.**
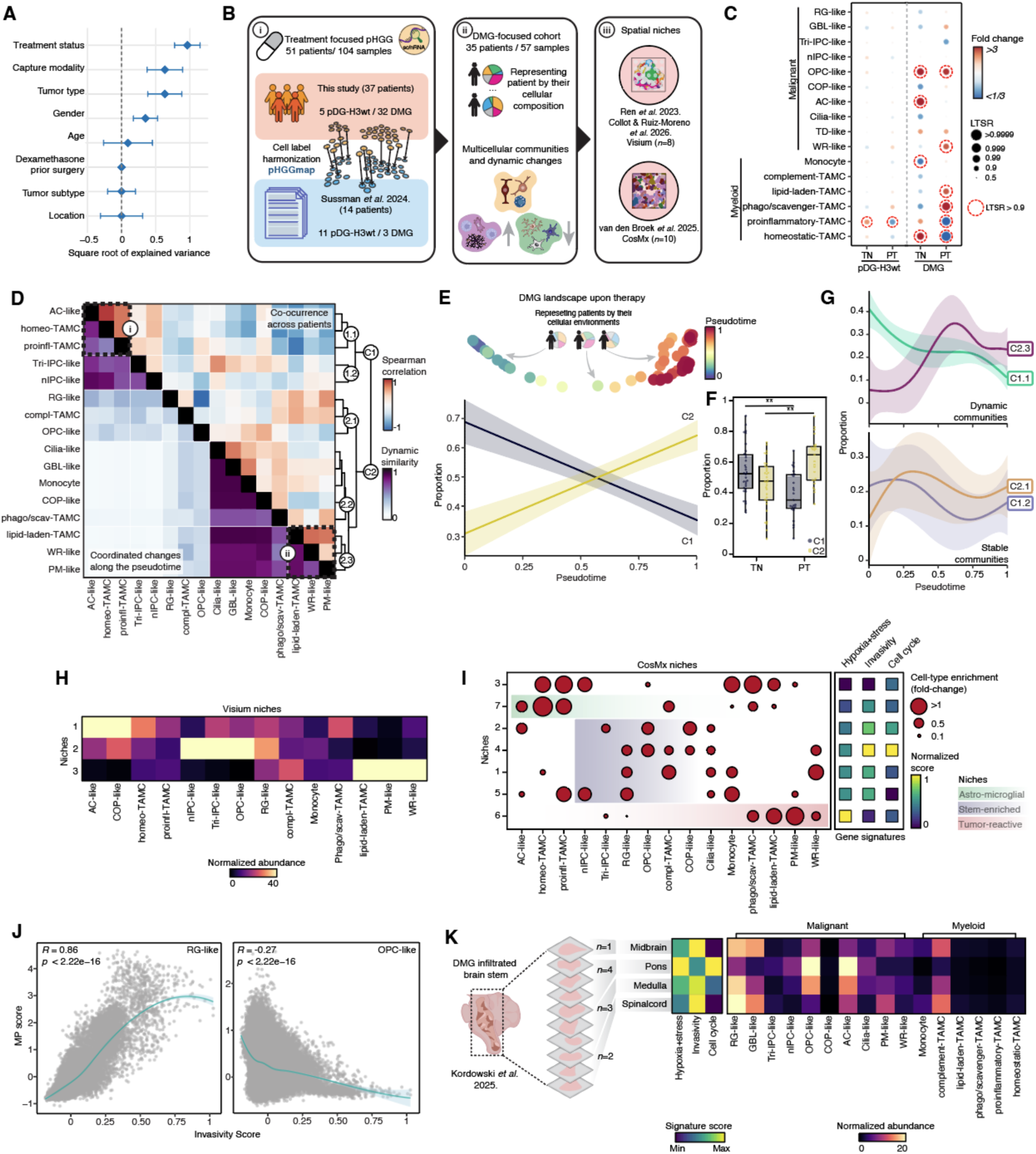
Therapy-induced shifts in multicellular communities/niches and invasive RG-like cells. (A) Forest plot showing the square root of the variance in cell type composition explained by each covariate and random effect using a linear mixed model^66,67^. (B) Schematic of the analytical pipeline. (C) Analysis of cell type proportions presents log_2_-transformed fold changes and LTSR scores, illustrating differences in the abundance of cell clusters based on treatment status and tumor type. The LTSR ranges from 0 to 1, where a value greater than 0.9 indicates a reliable estimate. TN: treatment naïve, PT: post-treatment. (D) Heatmap of pairwise correlations of proportions and cellular dynamics. Co-occurrence is shown at the top and similarity in dynamics at the bottom, with cell states grouped into communities. (E) PHATE embedding of DMG patients colored by pseudotime. Each dot represents one sample. Bottom, dynamic community patterns observed along the DMG inferred trajectory. (F) Box plot of the relative contribution of detected communities stratified by clinical status. Statistical significance between groups was assessed using a two-sided Mann-Whitney U test. \*\**p* < 0.01. (G) Dynamics of selected subcommunities’ proportions over pseudotime. The error bands represent the 95% confidence intervals. (H) Heatmap showing the normalized distribution of estimated cell type abundance as a percentage across the detected Visium (*n*=8) niches. (I) Cell-type enrichment within distinct spatial niches (CosMx, *n*=23), determined by cell state enrichment and tumor area. Normalized scores for selected signatures are shown. (J) Pearson correlation of RG- and OPC-like and invasivity signature^71^ scores in DMG malignant cells. (K) Left, heatmap depicts the signature scores per tumor region. Right, it shows the normalized distribution of estimated cell type abundance as a percentage across different areas for the multisector dataset from Kordowski *et al.*^73^ See also **Figure S5**.

### Therapy-associated shift yet persistent multicellular niches in DMG

Following our observation that DMG exhibits therapy-related changes, we focused our analysis on this tumor subtype (202,793 cells from 35 patients across 57 samples) (**Figure 5B-ii**). We previously identified two spatially distinct niches: one associated with tumor-reactive cancer cells (formerly referred to as MES-like), enriched in MDMs; and another consisting of AC-, OPC-, and COP-like (previously referred to as OC-like) cancer cells (AOO region), along with homeostatic MG, with AC-like cells displaying the closest spatial connection to this myeloid cell type^52^. We therefore initially re-evaluated these two niches using our updated classification of both cancer and myeloid cell populations. This enabled us to redefine how cell states co-occur across patients, referred to as multicellular communities in the DMG-treatment-focused cohort, and assess whether their composition shifts in response to treatment. To achieve this, we first normalized each subtype to obtain relative abundances. We then computed Spearman correlations of these abundances across all samples and represented each patient’s cellular landscape in a high-dimensional manifold, as previously implemented^70^ (**Figure 5B-ii**). This analysis revealed two major multicellular communities (**Figure 5D**): Community 1 (C1), predominant in treatment-naïve samples (median treatment-naïve = 0.52; post-treatment = 0.35), and Community 2 (C2), significantly enriched in post-treatment samples (median treatment-naïve = 0.48; post-treatment = 0.65), yet both communities were present in most samples **(Figures 5E and 5F**). Therapy-induced changes primarily affected cells in opposite ways at both ends of the C1/C2 axis (**Figures 5E-G, and S5C-E**): C1.1 subcommunity composed of AC-like cancer cells and MG-enriched TAMC programs (**Figure 5D-i**), and C2.3 subcommunity encompassing PM-, WR-like cancer programs and lipid-laden TAMCs (**Figure 5D-ii**). This analysis refined the composition of the previously defined MES region, resolving it into PM- and WR-like cancer programs together with lipid-laden TAMCs (*tumor-reactive*). It further confirmed the close association between AC-like cells and homeostatic- and proinflammatory-like MG (*astro-microglial*). To assess the spatial co-occurrence of these immune and cancer cell subsets, we reanalyzed our previous and publicly available DMG Visium ST datasets^19,52^ and single-cell resolution CosMx dataset^65^ (**Figure 5B-iii**). Among other niches, both the astro-microglial and tumor-reactive multicellular communities were consistently colocalized within spatial domains (Visium niches 1 and 3; CosMx niches 7 and 6, respectively, **Figures 5H and 5I; Table S7**), confirming their robustness and stable spatial organization in DMG. Importantly, this first layer of organization is associated with environmental factors, where the tumor-reactive niche is expectedly found in regions with high hypoxia and stress (niche 6), whereas the astro-microglial (niche 7) occupies areas characterized by lower levels (**Figure 5I and S5F**). Thus, in this study, we further demonstrate that despite treatment-induced shifts in their relative abundances, both niches persist across patient samples (**Figure 5G**, dynamic communities), underscoring their role as fundamental microenvironments in DMG.

### Invasive RG-like cells

Another notable microenvironment revealed by analysis of the spatial data was a domain highly dominated by stem/progenitor-like cancer cells (*stem-enriched*), including nIPC-, Tri-IPC-, OPC-, and RG-like populations (Visium niche 2, **Figure 5H**). Our Spearman correlation analysis in the DMG-treatment-focused cohort indicated that these stem-like populations, corresponding to subcommunities C1.2 (nIPC- and Tri-IPC-enriched) and C2.1 (OPC- and RG-enriched), underwent minimal changes in proportion in response to treatment (**Figure 5G**, stable communities), indicating that these cellular states remain largely preserved during the course of therapy. This stability raises the possibility that stem/progenitor-like-dominant multicellular communities may serve as reservoirs for therapy-resistant cells, representing a critical microenvironmental target. Leveraging the higher resolution of the CosMx dataset^65^, we could further define five spatial niches (1-5) containing stem-like cancer cell populations (**Figure 5I**). Among them, niche 4, which comprised RG- and OPC-like cells (**Figure 5I)** and closely resembled the subcommunity C2.1 across snRNA-seq samples (**Figures 5D**), exhibited the highest levels of proliferation and invasiveness^71^. Importantly, further analysis revealed that the invasivity signature^71^ was primarily driven by the RG-like population rather than the OPC-like cells, as confirmed by both sc/nRNA-seq and ST data (*R* = 0.86, *p* < 2.2e-16; **Figures 5J, S5G and S5H**). RG-like cells have been an underappreciated cancer cell population in DMG, particularly in comparison to proliferative OPC-like cells, which are more commonly regarded as central drivers of disease pathogenesis^20^. However, recent studies have identified RG-enriched niches at the invasive fronts of tumors in both adult and pediatric high-grade gliomas^19,72^. Consistent with this, deconvolution of a single-DMG multisector spatial dataset spanning 10 tissue sections across brain regions from the midbrain to the spinal cord^73^, revealed an increase in RG-like cells in nearby spreading anatomical areas from the pons, the tumor’s region of origin (**Figures 5K and S5I; Table S7**). In contrast, most other cancer states did not show such a large-scale spatial-associated pattern. Interestingly, RG-like cells were distributed across multiple niches, suggesting a certain degree of functional and spatial versatility. At the upper end of the hypoxia-stress gradient, beside the tumor-reactive niche (niche 6), niches 1 and 5 were predominantly enriched in RG-like cells within the malignant compartment (**Figure 5I**), suggesting that RG-like phenotypes might also adapt in an environment-responsive manner. To further characterize the phenotypic diversity within this population, we leveraged hypoxia and stress enrichment patterns, delineating two distinct RG-like subpopulations in the discovery cohort (**Figure S5J**). The smaller fraction, representing 19.6% of RG-like cells, was hypoxia-adapted and enriched for gene ontology terms related to environmental stress responses. In contrast, the majority of RG-like cells, accounting for 80.4%, showed high invasiveness scores and were enriched for gene ontology terms related to high metabolic activity and neuron projection development (**Figures S5K and S5L**). Altogether, these findings highlight the adaptability of RG-like cells and a functional divergence within the RG-like compartment, with one subset influenced by local environmental cues, while a dominant infiltrative population likely drives tumor progression through heightened invasiveness.

### Plastic transition toward an RG-like state during DMG dissemination

A rare case of DMG with abdominal metastasis treated in our center presented a unique opportunity to investigate the possible role of RG-like cells in tumor dissemination. The patient developed hydrocephalus requiring a ventriculoperitoneal shunt, which, though uncommon, can enable metastatic spread^74,75^. Fifty-six days after shunt placement, the patient developed ascites and abdominal metastasis. We performed longitudinal transcriptomic and epigenetic profiling of two distinct regions of the primary tumor at diagnosis, peritoneal ascitic fluid, and peripheral blood (*n* = 54,050 cells; **Figure 6A**). The primary tumor consisted mainly of AC-like cells in the pons and OPC-like cells in the region close to the cerebellar peduncle, with no RG-like populations detected (**Figures 6B and S6A**). By contrast, the metastatic cells in the ascitic fluid were almost exclusively composed of RG-like infiltrative phenotype (**Figures 6C, S6B and S6C**). iCNV analysis revealed distinct genotypes among primary states and additional genomic alterations in metastatic RG-like cells (**Figure S6D**), suggesting clonal diversification and acquisition of features that may support survival and dissemination in peripheral environments. Transcriptomic profiling showed that the metastatic cells activated the cell cycle and underwent a metabolic shift, marked by upregulation of *MYC* and its targets, glycolysis and oxidative phosphorylation (**Figures S6E and S6F**). Importantly, despite lacking RG-like gene expression, chromatin accessibility showed that both AC- and OPC-like cells in the primary tumor exhibited accessible chromatin at RG-associated loci, suggesting a latent epigenetic readiness for transition (**Figure S6G**). Trajectory analyses revealed a potential transition primarily from AC-like rather than OPC-like toward RG-like in ascites: PAGA analysis showed strong connectivity between these states (**Figure 6D**), and Palantir-based inference revealed a shift from AC-like (*AQP4, CLU*) toward RG-like phenotype (*HES1, FABP7*) (**Figure 6E**). Motif enrichment analysis further revealed that SOX and AP-1 TF families (e.g., JUN, FOS, FOSL1/2, JDP2), the latter recently implicated in governing transitions among cancer states in GB^76^, were enriched in metastatic RG- and AC-like cells but not in OPC-like cells (**Figure 6F and 6G**), suggesting a preferential AC-to-RG-like transition. Collectively, our findings indicate a plastic RG-like transition that may fuel DMG spread, emphasizing targeting RG-like cells and their potential precursor(s) (e.g., AC-like) as a therapeutic vulnerability.

**Figure 6.**
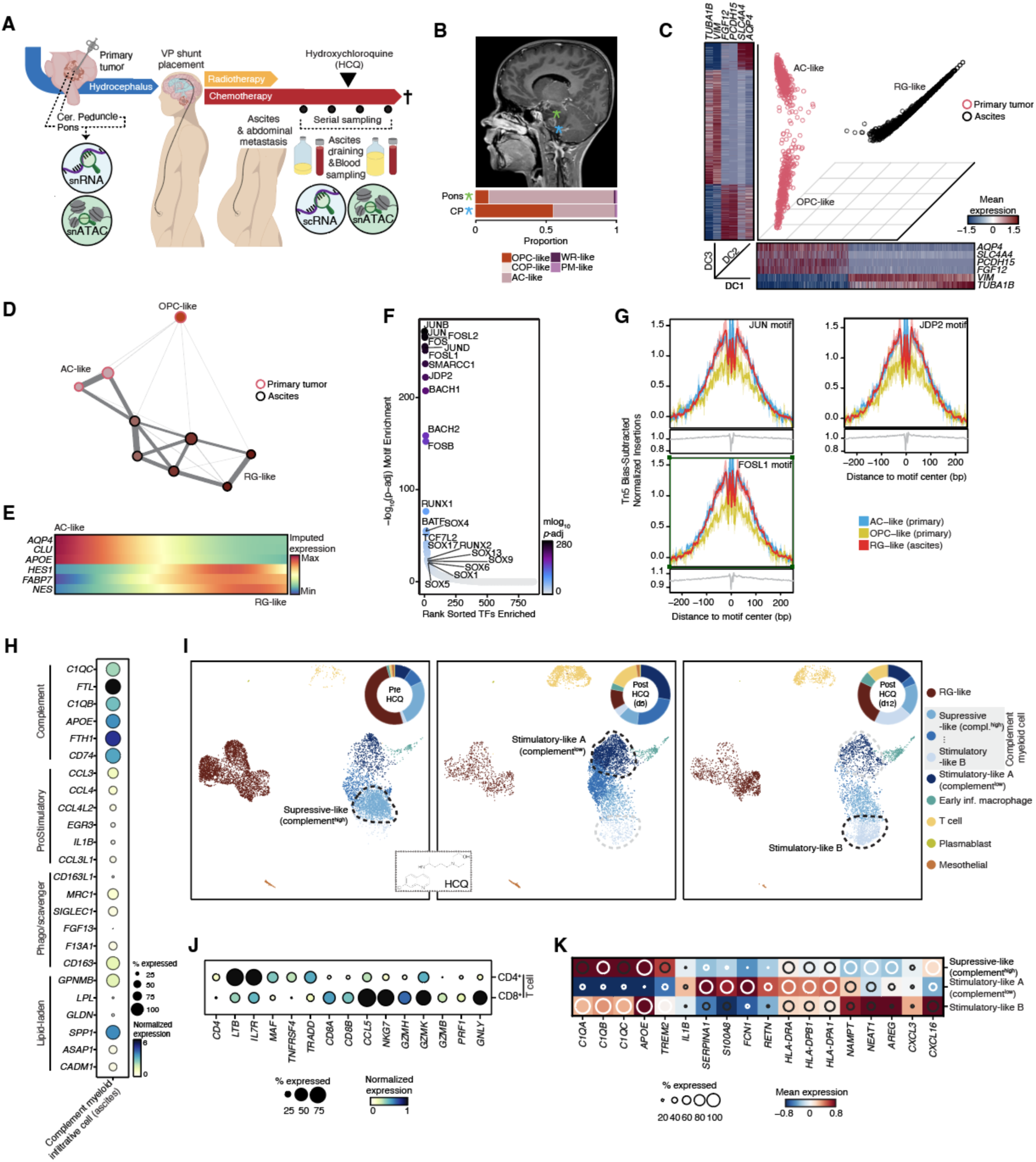
Adaptive plasticity, metastatic potential of RG-like cancer cells and TAMCs reprogramming in DMG. (A) Scheme of the clinical course and molecular tests of the patient with DMG and abdominal metastasis. VP: ventriculoperitoneal. (B) Sagittal and transverse MRI T1-enhanced plane. Stars mark the regions where biopsies were taken. Bottom, stacked barplot of the cell type proportion from the profiled tumor regions. CP: Cerebellar peduncle. (C) Diffusion map (DC1-DC2-DC3) of malignant cells from the primary tumor and ascites, colored by cancer phenotype. Heatmaps indicate mean gene expression along DCs. (D) PAGA graph^98^ of the primary and metastatic (ascites) cancer cells. Weighted edges indicate the strength of connectivity between clusters. (E) Gene expression trends along the AC to RG-like path. (F) Rank-sorted TF motifs colored according to the significance of their enrichment (-log10 adjusted *p*-value). Selected top-enriched TF motifs are labeled. (G) Tn5 bias-adjusted footprints for selected TFs in the malignant cells (primary and ascites). (H) Dot plot of gene expression of selected markers that define each immunomodulatory MP in the peritoneal infiltrative myeloid cells. (I) UMAP representation of the cells in the ascitic fluid collected at different time points before and after hydroxychloroquine treatment. Donut plot displays cell type proportions. HCQ: hydroxychloroquine. (J) Expression of gene markers that define T-cell subsets. (K) Expression of gene markers that define peritoneal infiltrative myeloid subsets. See also Figure S6.

### An RG-like-Complement-TAMC axis

Alongside the abundant metastatic RG-like cells, we observed a substantial infiltration of myeloid cells in the ascitic fluid. By concomitantly profiling PBMCs, we could follow the evolution of the peritoneal infiltrating myeloid cells from a CD14^+^ monocyte towards an early infiltrating macrophage, followed by the acquisition of an immunosuppressive phenotype (**Figure S6H**) that overlaps with our complement-TAMCs anti-inflammatory program in the primary tumor, as well as with the TAM C1QC phenotype defined in an immune pan-cancer atlas^56^ (**Figures 6H and S6I**). Interestingly, this finding correlates with the consistent association between RG-like cells and complement-TAMCs in our DMG-treatment-focused cohort (C2.1, **Figure 5D**) and spatial data (CosMx niches 1 and 4, **Figure 5I**). Our re-evaluation of sequential brainstem regions from the multisector spatial data^73^ indicated that, in addition to the infiltration of RG-like cells beyond the tumor’s core, there was also a progressive and concomitant increase in complement-TAMCs (**Figure 5K**). Furthermore, we observed a substantial overlap (56%, Jaccard = 0.39, FDR = 6.094746e^-26^) between the core gene programs of complement- and RG-like MPs (**Figure S6J**). It encompasses genes involved in antioxidant defense (*FTL, FTH1, GPX4, PRDX1*), cytoskeletal organization and motility (*VIM, ACTB, TUBA1B, CFL1*), and proteostasis regulation (*HSP90AB1, UBB*). It also includes secreted mediators such as *MIF*, which contributes to immune modulation and tissue remodeling^77^, and *APOE*, a lipid transporter that regulates complement activation and macrophage-driven neuroinflammatory signaling^78,79^ (**Figure S6K**). These transcriptional features were largely recapitulated in metastasis-associated RG-like cells and peritoneal complement-like macrophages (**Figure S6K**). Collectively, these observations point to a cooperative axis between RG-like and complement-TAMCs, in which reciprocal stress-, motility-, and immune-related signaling enhances cellular adaptability and may facilitate diffuse tumor invasion.

Previous studies demonstrate the plasticity and reversibility of TAMC programs, highlighting the potential to reprogram these cells into an anti-tumor phenotype^80–82^. Drawing on this possibility, the patient was treated with hydroxychloroquine as an adjuvant therapy in an attempt to repolarize the infiltrating macrophages and counteract their immunosuppressive nature^83–86^. This intervention led to a marked reduction in peritoneal RG-like cells, peaking at 5 days post-treatment, and was accompanied by increased recruitment of especially CD8⁺ T cells with a cytotoxic profile (e.g., *GZMB, PRF1, NKG7, GLNY*) within the ascitic compartment (**Figures 6H and 6I**). Although the number of complement-infiltrative peritoneal myeloid cells did not change drastically after treatment, subclustering identified various phenotypes. Some cells acquired an immunostimulatory signature (e.g., *IL1B*^hi^) after adjuvant therapy, marked by downregulation of complement and immunosuppressive markers like *APOE* and *TREM2* (stimulatory-like A). Another phenotype kept a complement-like profile but showed increased expression of chemotactic factors such as *CXCL16* (stimulatory-like B) (**Figure 6I, K)**. Both phenotypes exhibited higher levels of MHC-II genes and pathways involved in chemotaxis and antigen presentation (**Figures 6K and S6L)**. These results support that complement-TAMC might represent a moldable hybrid immune state, characterized by a predominantly immunosuppressive profile together with MHC-II expression and inflammatory features, suggestive of context-dependent adaptability within these myeloid cells. This flexibility is further illustrated by the increase of an immunostimulatory phenotype following hydroxychloroquine exposure, accompanied by enhanced CD8⁺ T cell recruitment and activation, and targeting peritoneal RG-like cells.

## DISCUSSION

Our study leverages single-cell multimodal profiling, integrating transcriptomics, epigenomics, and spatial analyses, alongside the latest human brain developmental and immune atlases, to resolve the complex cellular architecture of pHGG. This enabled the identification of diverse progenitor and differentiated phenotypes, as well as tumor-reactive states, ultimately defining ten distinct cancer cell programs. For instance, we identified a PM-like cancer state characterized by an (non-neuroglial) early developmental signature not evident in adult glioma, distinguishing pHGG from adult disease. Meanwhile, based on cell ontology and immunomodulatory states, we identified six key meta-programs (MPs) that characterize the dominant myeloid compartment and revealed how cancer and immune compartments interact within organized cellular communities. This large-scale, high-resolution, and spatial map uncovered a new therapeutic vulnerability for pHGG based on a specific cancer-immune state community.

Growing data suggest glioma cells appear to leverage phenotypic plasticity to adapt to fluctuating microenvironmental conditions, transitioning between states as needed^41,76^. Our findings are consistent with this model; we identified a WR-like program that is likely induced by environmental stressors, such as hypoxia. Epigenetic plasticity analysis revealed that these cells retain chromatin accessibility signatures from multiple developmental lineages, suggesting that diverse cancer states may converge to this phenotype in response to stress. This supports Mossi Albiach *et al.*, who reframed the MES-like phenotype in GB as a hypoxia-driven wound response that promotes shifts from developmental to WR-like program^41^. Such plasticity echoes models of transitions from stem-like or AC-like states to WR(“MES”)-like phenotypes^87,88^ and parallels injury-induced reprogramming, such as OPCs or oligodendrocytes converting into reactive astrocytes^89,90^. This mirrors behavior observed in adult high-grade glioma and reveals that such plasticity might also occur in pediatric brain tumors, highlighting a previously underappreciated capacity for dynamic state transitions in this context.

In line with this, our findings suggest that DMG cells may access a latent RG-like state, mimicking behavior reported in GB^91^, which could confer high invasiveness, enabling infiltration into distant regions and adaptation to diverse microenvironments. The plasticity of transition between states may be enabled or constrained depending on the cellular epigenetic landscape, as recently discovered in adult brain malignancies^92^. In our atypical DMG case where cancer cells metastasized into the abdomen, the AC-like phenotype, rather than the OPC-like state, was found to be more likely to transition into the invasive RG-like state. This observation may be linked to the fact that RG-like cells express several marker genes also found in AC-like cells^12^. It also aligns with earlier studies indicating a possible bidirectional transformation between radial glia and astrocytes^93,94^, implying that AC-like cancer cells might more readily reactivate an early developmental RG-like program. Furthermore, studies in both adult and pediatric gliomas have shown that RG-like cells, while transcriptionally aligned with astrocytic signatures, also express markers associated with WR programs^19,41^. This mixed expression profile has led to their classification as an intermediate state along the “reactive-developmental program” axis. We refined this ‘intermediate’ state by identifying two distinct subtypes of RG-like cells: one exhibiting a hypoxia-associated profile, and another with an infiltrative signature. The latter subtype, characterized by elevated metabolic activity and marked invasiveness, was detected in distant infiltrative regions by multisector spatial analysis and in the patient’s ascitic fluid with abdominal metastasis, supporting its role in tumor dissemination. Thus, RG-like state appears to retain developmental migratory programs, driving the infiltrative behavior characteristic of DMG and reflecting an inherently plastic, embryonic epigenetic state that enables adaptation to stress and opportunistic invasion.

Targeting RG-like and other high-risk phenotypes in diffuse glioma requires a deeper understanding of their microenvironment. In particular, interactions with the dominant myeloid compartment in pHGG may uncover new immunotherapeutic strategies. We identified three predominant, conserved multicellular communities (sc/nRNA level) and spatial niches (ST level). One niche forms a tumor-reactive ecosystem that is prominently present in post-treatment samples. It includes PM- and WR-like tumor cells together with lipid-laden TAMCs, all exhibiting a shared transcriptional response to environmental cues such as hypoxia. In contrast, the second niche, more predominant in treatment-naïve samples, contains AC-like tumor cells and homeostatic or pro-inflammatory MG, forming a more differentiated and immune-inert, neuroglial-like microenvironment. Importantly, we discovered that RG-like tumor cells were spatially associated with and shared a gene signature with complement-TAMCs. This niche persists across inferred treatment trajectory, further underscoring its role not only in tumor progression but also potentially in treatment resistance. Altogether, these three cancer-dominant niches represent core functional units of pHGG biology and, importantly, involve distinct myeloid populations. In adult GB, current efforts aim to reprogram lipid-laden macrophages^63^ to enhance antitumor responses. In our recent work, these cells were also identified in pHGG^65^, where their targeting will likely modulate the tumor-reactive niche. In contrast, our previous research in DMG identified the IGSF11-VISTA axis^52^ as a key mechanism for reprogramming homeostatic MG into anti-tumor or phagocytic states, targeting the astro-microglial niche. Building on this, we now show that complement-TAMCs may serve as a promising target for reprogramming efforts aimed at targeting the highly invasive RG-like cancer cell state, an essential member of the stem-enriched niche. We presented a pioneering case study involving the treatment of a patient with abdominal metastasis using hydroxychloroquine. This was associated with increased immunostimulatory features in complement-TAMCs and promoted the recruitment of cytotoxic CD8^+^ T cells and an antitumor response against RG-like cells in the ascites. However, further investigation is needed to develop more specific therapeutic strategies for targeting complement-TAMCs, beyond broadly acting agents, such as hydroxychloroquine, and to confirm that this can reduce tumor burden via targeting its most invasive features.

Altogether, our work highlights a central challenge in designing effective immunotherapeutic strategies for pHGG: the potential need to simultaneously modulate distinct myeloid populations that occupy functionally and spatially segregated niches. Lipid-laden TAMCs, homeostatic and pro-inflammatory MG, and complement-expressing TAMCs each preferentially engage with distinct cancer cell states and respond to different environmental cues. As such, rewiring these heterogeneous myeloid phenotypes will likely require combinatorial or highly selective approaches tailored to the specific biology of each niche. Achieving this level of precision remains a major hurdle, but also a critical opportunity, to unlock sustained antitumor responses in pHGG. We anticipate that the outcome of our research, the pHGGmap, will serve as a comprehensive reference to explore myeloid-cancer cell interactions, guide therapeutic target discovery, and inform the rational design of next-generation immunotherapies for pediatric high-grade glioma.

## Limitations of the study

Our study has several limitations. First, cellular “ecotypes” and state proportions vary across tumor regions. Most patients were sampled at a single site and time point, so our pHGGmap may overgeneralize local features to the whole tumor. Second, the inferred AC-to-RG-like transition during dissemination is supported by correlative transcriptomic/epigenomic evidence but not proven causally. We cannot exclude alternative explanations, such as the selection of pre-existing RG-like cells outside the CNS, or that multiple phenotypes seed metastases, but RG-like cells preferentially persist. Given that intracranial dissemination is more common, dissecting programs that enable spread within the CNS remains a priority. Third, therapy-associated shifts were assessed across heterogeneous regimens and variable biopsy timing; we lacked adequate numbers per protocol to attribute changes to specific treatments or to adjust for confounders such as steroid exposure or extent of resection. It is essential to recognize that most post-therapy samples analyzed in our study were derived from autopsy material, reflecting end-stage disease after extensive treatment and terminal progression. The observed shifts, therefore, likely represent a dynamic process, the precise timing of which remains unresolved. Fourth, despite our efforts to mitigate the impact of technical factors, remaining technical biases could still affect state calling and composition estimates, as factors like capture modality (cells vs. nuclei, discussed further in **Technical note**), platform/library differences, dissociation bias, batch correction/integration choices, and spatial platform resolution/panel content all add variability. Finally, while tumor-myeloid communities appear robust before and after therapy, our ability to assess their prognostic significance was limited. The relatively short survival time of most patients and the small number of cases with longitudinal sampling reduced the statistical power to test whether specific community configurations predict treatment response, recurrence timing, or overall survival. Prospective multi-region, longitudinal sampling with standardized clinical metadata, responder/non-responder cohorts, and experimental perturbations will be required to determine how/when/why cell transitions occur and establish causality and therapeutic leverage of the RG-like-complement axis.

---

**Figure S1.**
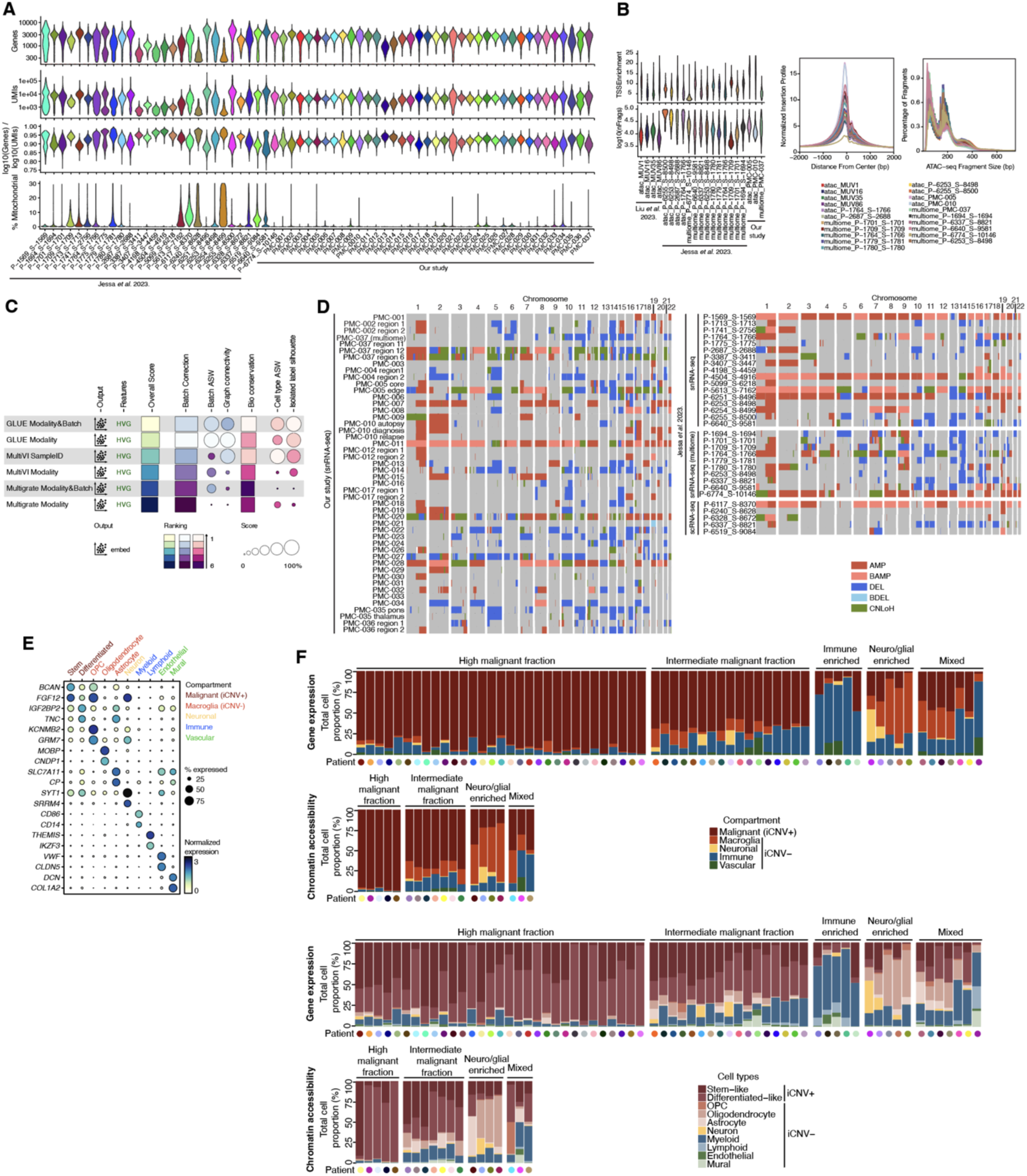
Construction of a multimodal map of pHGG, related to Figure 1. (A) Violin plots show the distribution of the number of genes detected, UMI counts per nucleus, cell complexity (log_10_(genes) / log_10_(UMIs)), and mitochondria percentage for each patient (discovery cohort) after stringent quality filtering. (BViolin plots (left) show the distribution of the transcription start sites (TSS) detected and the number of fragments per nucleus (log_10_ transformed). Right, TSS enrichment profiles and fragment size distributions for each patient (discovery cohort). (C) scIB benchmarking^27^ of three different methods for mosaic single-cell multimodal integration. (D) Heatmap of the consensus copy number segments per sample inferred from the sc/nRNA-seq. (E) Dot plot of gene expression for selected markers that define each cell type. (F) Stacked bar graphs showing the proportion of cells per compartment in the discovery cohort (sc/nRNA-seq and snATAC-seq). Patients were classed into five categories according to the enrichment of a particular cell compartment/cell type. Colored dots (patient) matched the color scheme in Figure 1C.

**Figure S2.**
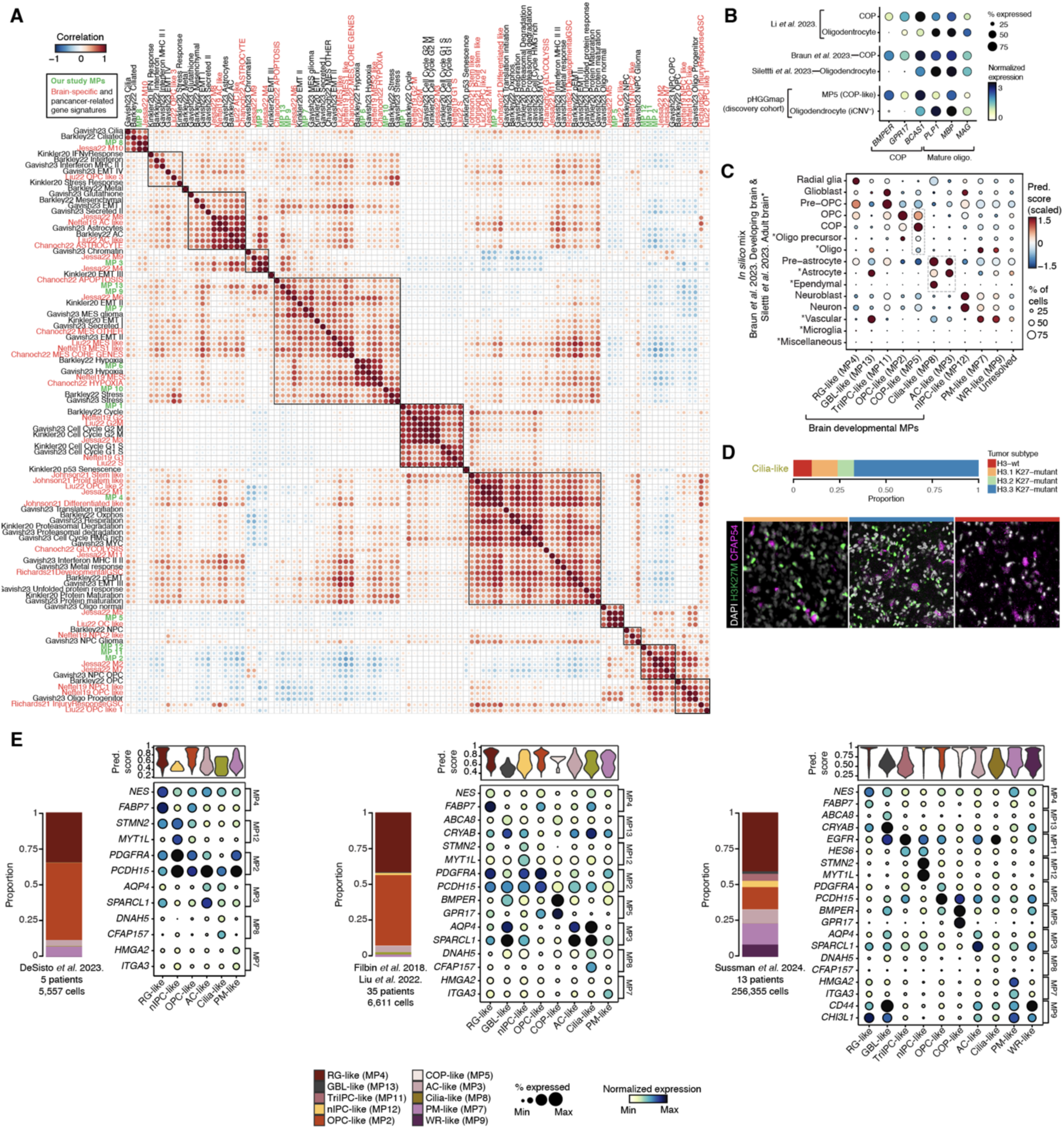
Characterization of developmental molecular states in malignant pHGG cells, related to Figure 2. (A) Correlation matrix depicting the expression signatures of NMF MPs compared with those specific to DMG and other signatures related to brain tumors or pan-cancer across published studies^17,20,30–36^. (B) Dot plot of the expression of classical COP and mature oligodendrocytes markers in healthy references and pHGGmap. (C) Dot plot displays prediction scores of pHGG cancer cells (per MP) after mapping *onto in silico* mix of reference atlases^22,37^. Original annotation per reference was kept. (D) Stacked bar graph showing the proportion of cilia-like cells present in different tumor subtypes (sc/nRNA pHGGmap). Immunofluorescence validation (bottom) using the cilia-like marker CFAP54 (magenta). (E) Stacked bar graphs show the proportion of MPs per study. Dot plots show the expression of key markers that define our cancer MPs for the Filbin *et al.*^15^, Liu *et al.*^20^ (Smart-seq2), DeSisto *et al.*^25^ (10X), and Sussman *et al.*^18^ (10X) datasets (validation cohort). On top, the violin plots represent the prediction scores per MP.

**Figure S3.**
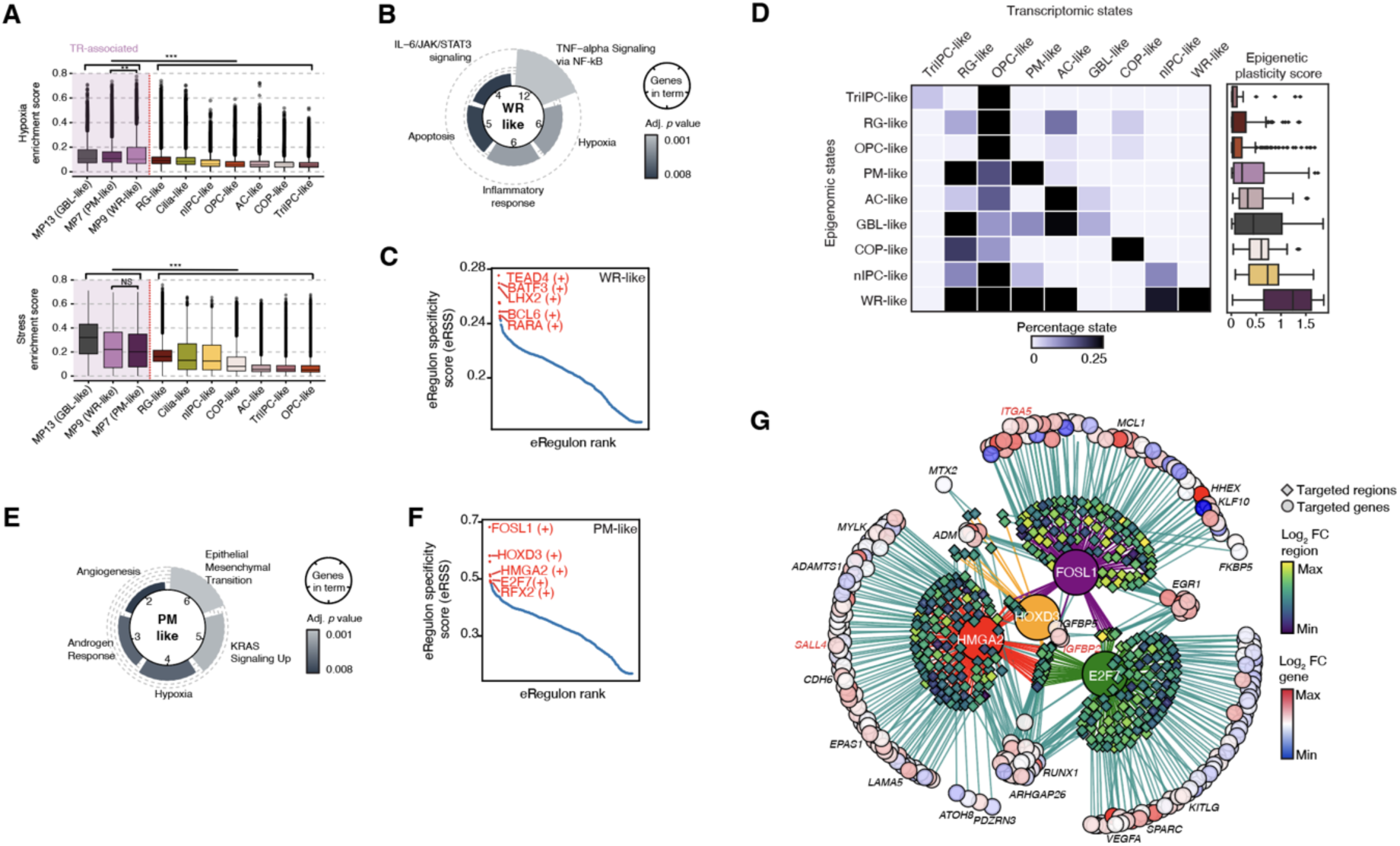
Transcriptomic and epigenomic characterization of tumor-reactive states in malignant pHGG cells, related to Figure 3. (A) Box plots show the enrichment score of the hypoxia and stress MPs for each developmental and tumor-reactive-related MP. Ordered by their average mean, from highest to lowest. Statistical significance between MP7 and MP9 was assessed using a Wilcoxon test. A Welch’s t-test was applied to compare the enrichment scores of the top three MPs against the scores of the remaining MPs. \*\**p* < 0.01, \*\*\**p* < 0.001. (B) Circular diagrams display the top 5 enriched HALLMARKS. In the center, the number of genes associated with each term is indicated. Dark shades represent adjusted *p*-values. (C) Scatter plot displays the eRegulon specificity scores for each TF alongside their corresponding regulon for WR-like cells. The five highest-ranking TFs and eRegulons are marked in red. (D) Confusion matrix of classifiers illustrating the correspondence between epigenomic and transcriptomic cancer cell state classifications. Each entry represents the number of metacells from an epigenomic cluster assigned to a transcriptomic cluster, normalized by row to reflect relative cluster overlap. (E) Circular diagrams display the top 5 enriched HALLMARKS. In the center, the number of genes associated with each term is indicated. Dark shades represent adjusted *p*-values. (F) Scatter plot displays the eRegulon specificity scores for each TF alongside their corresponding regulon for PM-like cells. The five highest-ranking TFs and eRegulons are marked in red. (G) Visualization of the eGRN formed by HMGA2, HOXD3, FOSL1, and E2F7. Target nodes for transcription factors focus on genes and regions with high variability.

**Figure S4.**
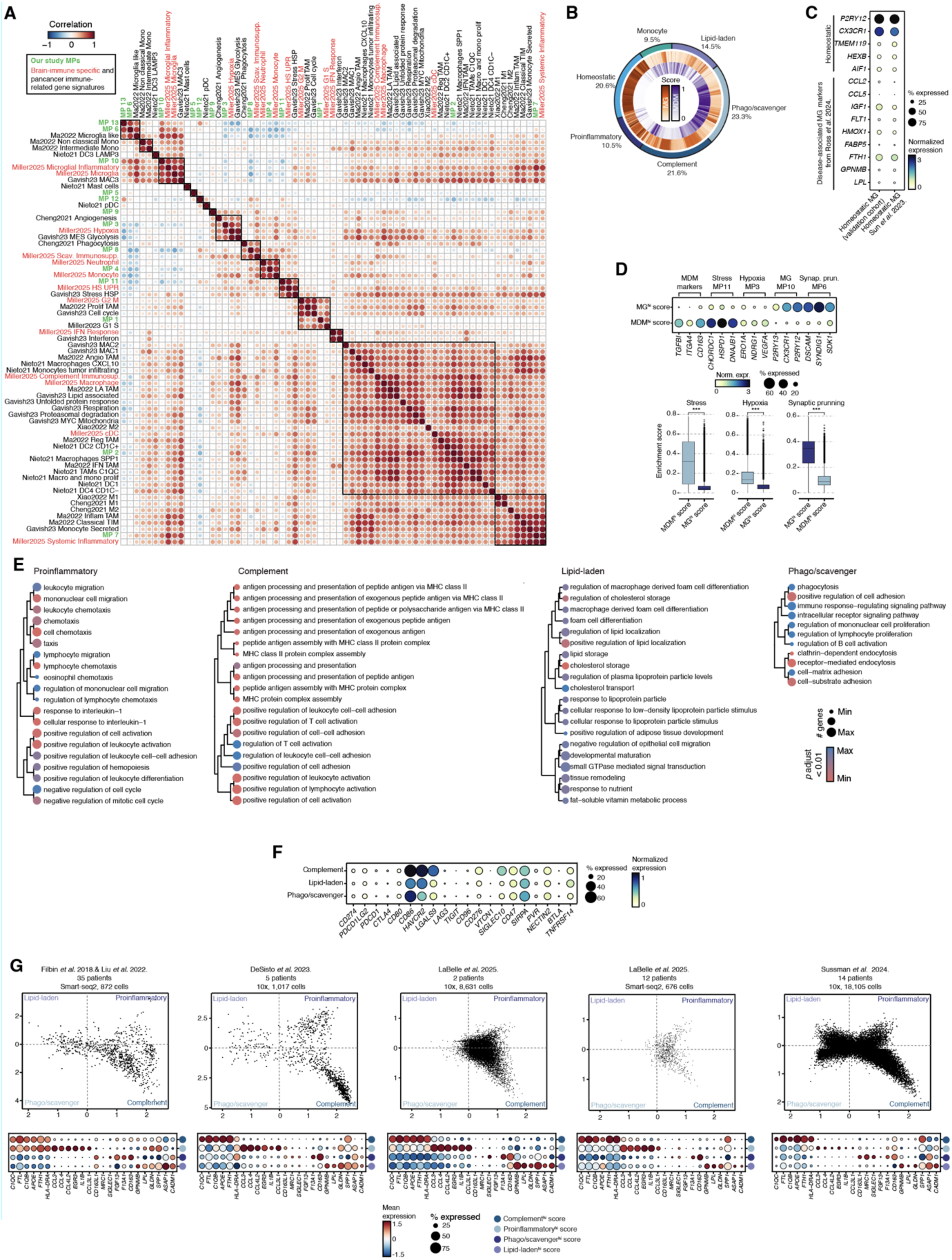
Transcriptomic characterization of molecular phenotypes of myeloid cells in pHGG, related to Figure 4. (A) Correlation matrix depicting the expression signatures of NMF MPs compared with those specific to myeloid cells and other signatures related to brain tumors or pan-cancer across published studies^14,30,53–56^. (B) Donut plot displays the enrichment for MDM and MG signatures and the proportion of immunomodulatory programs. (C) Expression of gene markers that define homeostatic MG and DAMG across cells derived from pHGGmap, and Sun, *et al.*^59^ (D) Dot plot of gene expression of selected markers that define each MP (top). Box plots (bottom) show the enrichment score of core biological MPs for MDM and MG. Statistical significance was assessed using a Wilcoxon test. \*\*\**p* < 0.001. (E) Dot plot of the top significant pathways per immunomodulatory MP from the GO:BP database. (F) Expression of immune checkpoint/inhibitory genes in immunosuppressive TAMCs. (G) States plot for TAMCs obtained from Filbin *et al*.^15^, Liu *et al*.^20^, DeSisto *et al*.^25^, LaBelle *et al*.^26^, and Sussman *et al.*^18^ (validation cohort). Every quadrant represents a distinct inflammatory TAMCs MP, and each dot is an individual cell. On the bottom, the dot plot displays the mean expression of selected markers that define each inflammatory MP per study.

**Figure S5.**
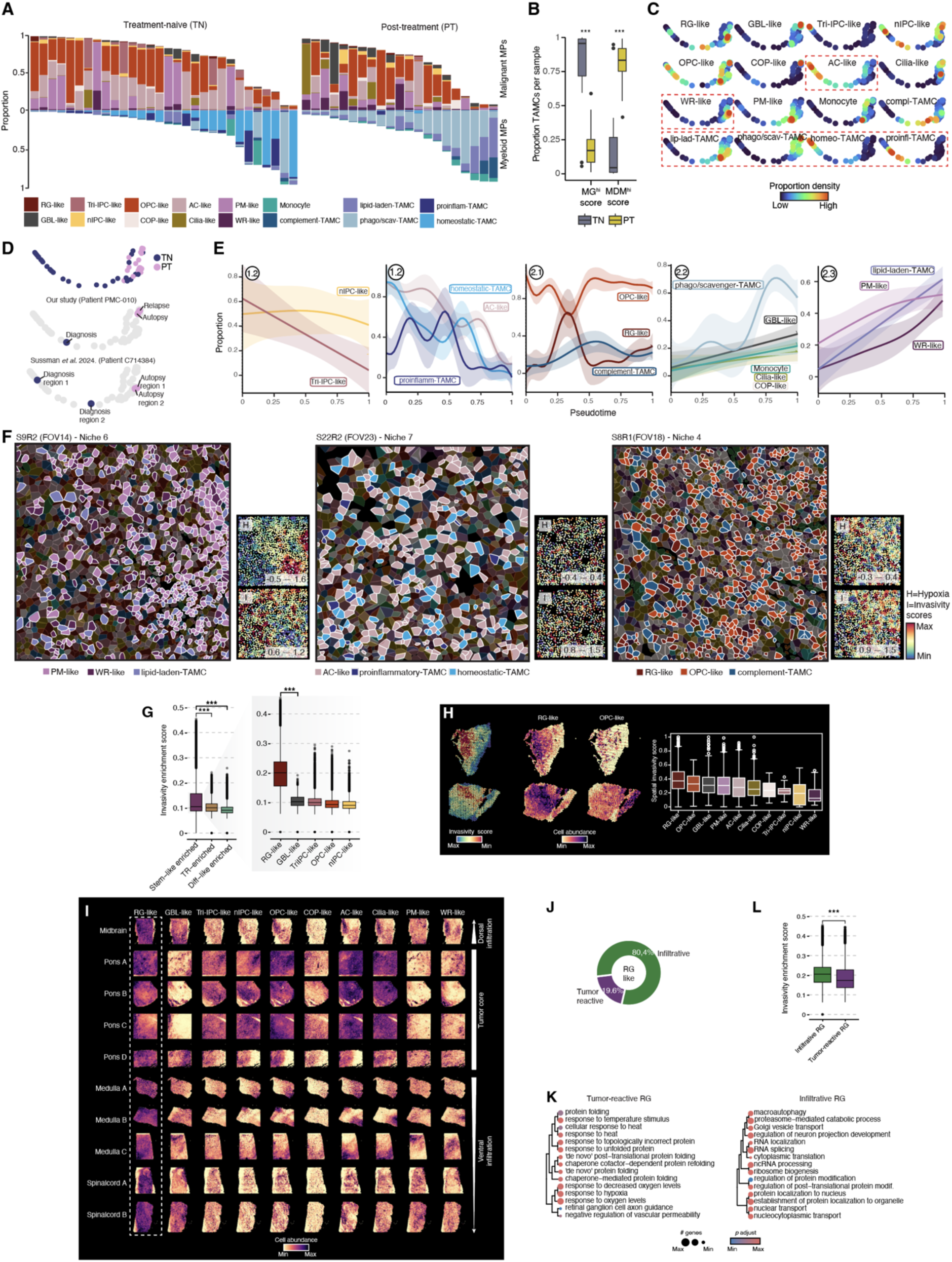
Dynamic changes in the DMG TME upon therapy and invasive features of RG-like cancer cells, related to Figure 5. (A) Barplot of the cell type proportions in pHGG patients split by clinical status. Only the malignant and myeloid MPs are displayed. (B) Box plots of the percentages of TAMCs (MDM and MG) compared between treatment-naïve (TN) and post-treatment (PT) tumors. Statistical significance was assessed using a Wilcoxon test. \*\*\**p* < 0.001. (C) Selected cell state proportions across the DMG PHATE embedding. (D) DMG PHATE embedding colored by treatment status. Selected patients with longitudinal sampling are shown. (E) Dynamics of cell state proportions over pseudotime in cellular subcommunities. The error bands represent the 95% confidence intervals. (F) Selected FOVs illustrate a range of DMG niches in the CosMx dataset. Segmented cells are colored based on the most likely predicted cell subtype. A white contour line outlines cell types that are enriched in each niche. Right of each niche, spatial gene expression signature scores. (G) Box plot shows the enrichment score of the invasivity signature for each malignant niche. Zoom-in shows cancer states in the stem-like niche, ordered by their average mean, from highest to lowest. Statistical significance between the first and the second highest was assessed using a Wilcoxon test. \*\*\**p* < 0.001. (H) Spatial invasivity signature^71^ score and RG- and OPC-like estimated cell type abundance. Box plot shows the invasivity score across estimates of spatial malignant MPs. (I) Estimated cell abundances for each cancer MP split by each tumor location from the Kordowski *et al.* dataset^73^. (J) Donut plot displays the proportions of RG-like subtypes. (K) Dot plot of the top significant pathways per RG-like subtype from the GO:BP database. (L) Box plot shows the enrichment score of the invasivity signature for each RG-like subtype. Statistical significance was assessed using a Wilcoxon test. \*\*\**p* < 0.001.

**Figure S6.**
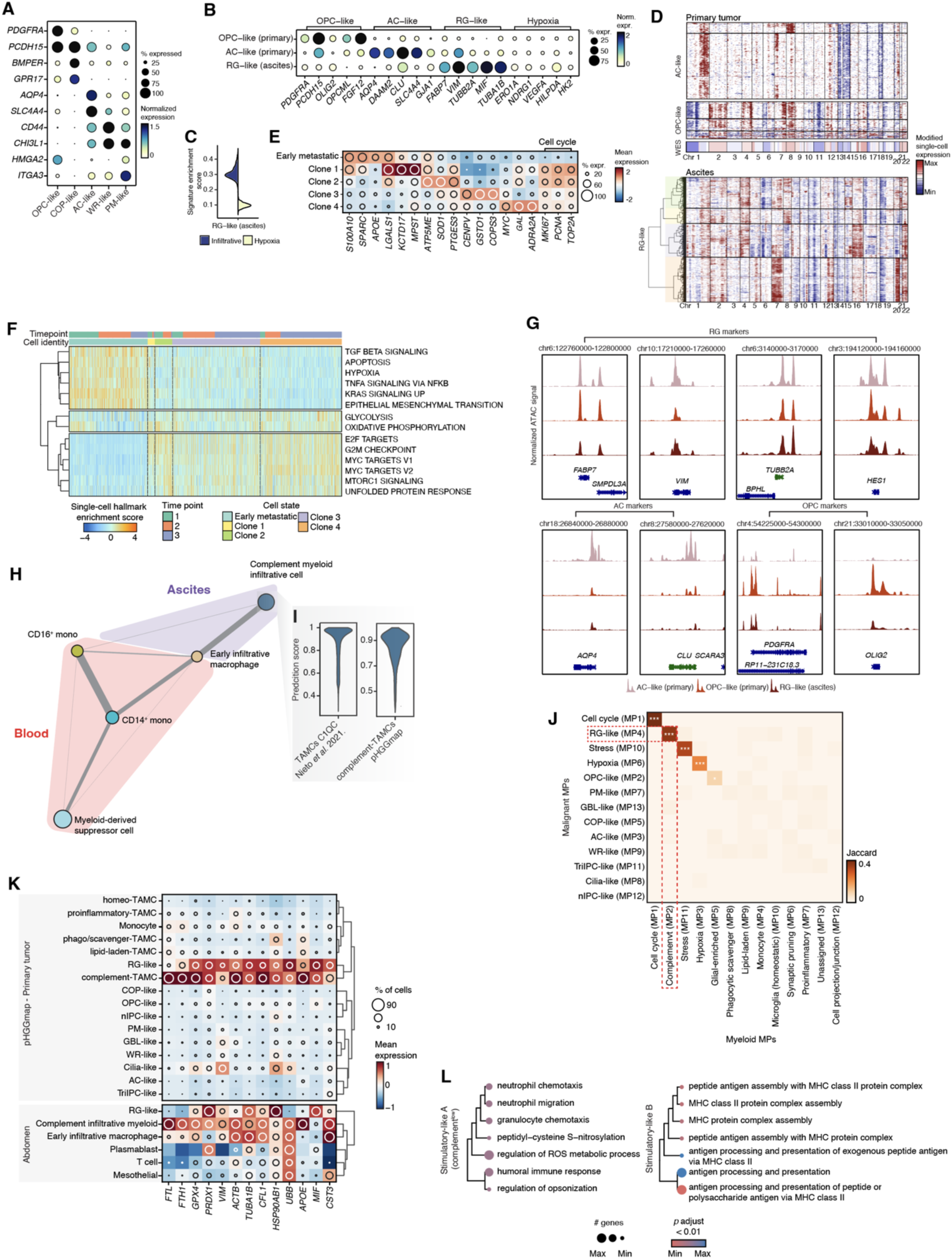
RG-like invasive and adaptive features and TAMCs modulation, related to Figure 6. (A) Expression of gene markers that define malignant phenotypes in the primary tumor. (B) Expression of gene markers that define AC- and OPC-like in the primary and RG-like in the ascitic fluid. (C) Violin plots depict the scores of selected gene signatures in the abdominal RG-like cells. (D) Clustered heatmap of iCNV profiles of primary and ascitic cancer cells clustered into genetic subclones based on CNV pattern. Bottom panel in primary iCNV: copy number profile derived from bulk exome sequencing data. (E) Expression of gene markers that define ascitic malignant cells. (F) Heatmap displays GSEA of individual metastatic cells using the HALLMARKS collection from the MSigDB. Cells are colored by enrichment score and hierarchically clustered within each cell identity and time point. (G) Genome tracks of aggregate scATAC-seq profiles around the selected genes per cancer state. (H) PAGA graph^73^ of PBMCs (blood) and infiltrating myeloid cells (ascites). Weighted edges indicate the strength of connectivity between clusters. (I) Violin plot displays the prediction score of the peritoneal infiltrative myeloid cells when performing reference mapping using Azimuth^97^. (J) Jaccard similarity heatmap illustrating the significant correspondence of genes between malignant and myeloid MPs. *p*-values were corrected for multiple testing using the Benjamini-Hochberg procedure. Adjusted values (FDR) are reported. *FDR < 0.05, ***FDR < 0.001. (K) Expression of selected shared gene markers between RG-like (MP4) and complement-TAMCs (MP2), separated by cells in the primary tumor (pHGGmap) and ascitic fluid (abdomen). (L) Dot plot of the top significant pathways (GO:BP database) of selected peritoneal infiltrative myeloid subsets.

## RESOURCE AVAILABILITY

### Lead contact

Requests for further information should be directed to and will be fulfilled by the lead contact, Hendrik G. Stunnenberg

### Materials availability

This study did not generate new, unique reagents.

### Data and code availability

- The newly generated snRNA-seq, snATAC-seq, and snMultiome raw sequencing data from this study will be available through controlled access in compliance with human privacy regulations. They will be obtained from the European Genome-phenome Archive (EGA) upon completion of a data use agreement (EGAXXXXXXX). Previously published datasets re-analyzed in the **P**ediatric-type diffuse **H**igh **G**rade **G**lio**MAP** (pHGGmap) were retrieved from the following repositories: EGAS00001005773, EGAS00001007035, GSE184357, GSE227983, GSE231860, https://data.humantumoratlas.org/publications/hta4_2024_biorxiv_jonathan-h-sussman, and https://singlecell.broadinstitute.org/single_cell/study/SCP147/single-cell-analysis-in-pediatric-midline-gliomas-with-histone-h3k27m-mutation.
- The pHGGmap discovery cohort (raw/normalized counts, cell-type annotations, and associated technical/clinical metadata) is publicly accessible via Zenodo (https://doi.org/10.5281/zenodo.17063631).
- Spatial transcriptomics datasets re-analyzed in this study are available at GSE194329, GSE280990, and https://zenodo.org/records/1422594.
- All code supporting this study will be deposited on GitHub (https://github.com/ccruizm/pHGGmap).
- Further information and requests for resources or data reanalysis are available from the lead contact upon reasonable request.

## ACKNOWLEDGMENTS

We extend our gratitude to the patients and their families for their participation in this research study. Appreciation also goes out to the clinicians, nurses, care teams, and pathologists at our institute with whom we collaborate closely, as well as the flow cytometry, imaging, and sequencing core facilities of the Princess Máxima Center. A special thank you is due to the patient, known as ‘The Flash’ (donor with DMG and abdominal metastasis) and his family for their contributions to this work. We acknowledge Marc van Wetering and Femke Ringnalda for their assistance with tissue acquisition and Julie Lammers for her help in sample preservation. Our gratitude extends to the high-performance computing resources in Utrecht and the Dutch national supercomputer Snellius for their support in computation and infrastructure. Additionally, we appreciate the meaningful discussions with Kimberly Siletti, Alejandro Mossi, and Piyush Kumar about the connections between cancer developmental-like programs and healthy brain development. C.R.M. especially thanks the European Joint Programme on Rare Diseases for the fellowship that allowed him to visit the Theis lab. This research was supported by the Foundation Children Cancer free (Kika) starting grant.

## AUTHOR CONTRIBUTIONS

Conceptualization - C.R.M., R.C., A.C.R., and H.G.S.; Data curation - C.R.M.; Formal analysis - C.R.M.; Funding acquisition - A.C.R., and H.G.S.; Investigation - C.R.M., R.C., G.D., and B.t.P.; Project administration - C.R.M., and H.G.S.; Resources - C.R.M., R.C., T.v.d.B, N.B., I.L.I., F.J.T., D.S.M., M.E.G.K., E.W.H., J.v.d.L., F.C., A.Z., E.H., and D.G.v.V.; Software - C.R.M.; Supervision - A.C.R. and H.G.S.; Validation - C.R.M. and R.C.; Visualization - C.R.M., R.C., A.C.R. and H.G.S.; Writing - C.R.M., R.C., E.J.W., A.C.R. and H.G.S.

## DECLARATION OF INTERESTS

F.J.T. consults for Immunai Inc., CytoReason Ltd, Cellarity, BioTuring Inc., and Genbio.AI Inc., and has an ownership interest in Dermagnostix GmbH and Cellarity. The other authors declare no conflicts of interest.

## METHOD DETAILS

### Human tissue collection

Surgical tumor samples (treatment-naïve) used in this study were retro- and prospectively selected from the biobank or collected after biopsy at the Princess Máxima Center for Pediatric Oncology. This research was reviewed and approved by the institutional biobank review committee (PMCLAB2020.142). The cohort comprised tumor tissues from pediatric patients who underwent surgical resection between 2018 and 2022. Informed consent for biobanking and the use of these tissues in research was obtained from patients and/or their legal guardians in compliance with national legislation and ethical guidelines stipulated by the Declaration of Helsinki (2013). Tissue diagnoses were based on comprehensive histological and molecular analyses, including standard histology, immunohistochemistry, EPIC methylation profiling, whole-exome sequencing, and whole-transcriptome sequencing.

Post-therapy and/or autopsy human tissue samples were acquired through established programs at the VU University Medical Center Diffuse Intrinsic Pontine Glioma Autopsy Program Caretti, 2013 #385 and the Tumor Donation Program at the Princess Máxima Center for Pediatric Oncology (Máxima-TDP). Both programs are approved by the Medical Research Ethics Committee NedMec (MEC 22-589/PB22TUM). Informed consent was obtained from parents and, when applicable, patients older than 12 years capable of providing consent, in accordance with national laws and the Declaration of Helsinki (2013). Brain biopsies and post-mortem tissues were collected through stereotactic tumor biopsy or brain/full-body autopsies, respectively. Samples were immediately flash-frozen, embedded in OCT compound (Sakura Finetek), and stored at -80°C until processing. Cryosections of 10 μm thickness were prepared using a LEICA Cryostar NX70 cryostat and utilized for downstream single-nucleus RNA sequencing (10x Genomics 3’ v3.1).

For the case involving abdominal metastasis of DMG, ascitic fluid was collected through sterile paracentesis. In cases of recurrent fluid accumulation, a palliative surgically placed abdominal drain facilitated long-term fluid collection. Samples were maintained in sterile collection systems. Ascitic fluid was immediately centrifuged at 300g for 10 minutes at room temperature and subsequently washed three times under the same conditions with tumor medium (Advanced DMEM/F12 supplemented with Penicillin-Streptomycin, GlutaMAX, and HEPES Buffer Solution; Gibco Laboratories). If recovered cells contained more than 30% red blood cells (RBCs), an RBC lysis solution (Miltenyi Biotec) was employed. Peripheral blood mononuclear cells (PBMCs) were isolated from fresh EDTA peripheral blood using density-gradient centrifugation (Lymphoprep; Stem Cell Technologies). Cells obtained from ascitic fluid and PBMCs were either immediately processed for single-cell RNA sequencing or placed in freezing medium (peritoneal cells: tumor medium with 10% DMSO; PBMCs: 40% RPMI Medium, 50% FBS, and 10% DMSO) and stored at -80°C or in liquid nitrogen until further use.

### sc/nRNA-seq sample and library preparation

Several 10 μm cryosections per specimen were collected into an ice-cold tube and mixed with 500 μL of RNase-inhibiting lysis buffer containing 0.1% NP40 in a Tris-based solution. The mixture was homogenized in a glass-on-glass dounce homogenizer on ice, using a loose pestle for 10 strokes and a tight pestle for 15 strokes. The nuclei solution was filtered through a 70 μm Flowmi cell strainer and centrifuged at 4°C at 500g for 5 minutes. The pellet was resuspended in 1.5 mL Wash Buffer (Tris-based, 2% BSA, RNase inhibitor) and filtered through a 40 μm Flowmi strainer. DAPI staining was applied for nuclei preparation, followed by sorting intact nuclei on a Sony SH800 cell sorter with a 100 μm nozzle. Approximately 15,000 nuclei per sample were sorted directly into 10x RT Buffer and processed per the Single Cell Gene Expression 3’ v3.1 protocol (10x Genomics). For scRNA-seq from ascitic fluid, fresh samples were washed with a warm medium (Advanced DMEM/F12, enriched with 10% FBS) and pelleted three times. The cryopreserved cells underwent thawing in a 37 °C water bath for two minutes, then washed with warm medium and centrifuged at 350g for 5 minutes, followed by three additional washes. Both fresh and cryopreserved cell pellets were resuspended in an appropriate volume of sterile PBS (Sigma-Aldrich) with 0.04% BSA (Sigma-Aldrich) to achieve a concentration of 1,200-1,500 cells/μl. Cell viability was evaluated using 0.1% Trypan blue staining and measured with a Countess II Automated Cell Counter (Thermo Fisher Scientific). If the viability of the ascitic cells (fresh or cryopreserved) was below 70%, the Dead Cell Removal Kit (Miltenyi Biotec) was utilized according to the manufacturer’s instructions. Approximately 15,000 cells per sample were processed per the Single Cell Gene Expression 3’ v3.1 protocol (10x Genomics). Library quality was assessed using a 2100 Bioanalyzer (Agilent) and sequenced on a NovaSeq 6000 (Illumina). The targeted sequencing depth per cell was 50K reads.

### snATAC-seq sample and library preparation

For single-nucleus ATAC sequencing (snATAC-seq), nuclei were isolated following the recommended protocol from 10x Genomics, using suggested reagents and consumables. Tissue-specific adjustments were made to the lysis buffer concentration and incubation times: primary tumors were incubated with 0.1X lysis buffer for 10 minutes on ice, whereas ascitic cells were incubated with 1X lysis buffer for 5 minutes on ice. Following isolation, nuclei were washed, pelleted, and resuspended in chilled 1X nuclei buffer (10x Genomics). Samples were filtered through a 40 µm Flowmi cell strainer (Bel-Art) to remove cellular debris and large aggregates. Nuclei counts and viability were determined using 0.1% trypan blue staining on disposable slides with a Countess II Automated Cell Counter (Thermo Fisher Scientific), while nuclear morphology and quality were visually assessed using a Zeiss light microscope. Approximately 10,000 nuclei per sample underwent upfront tagmentation with Tn5 enzyme following the Single ATAC v1.1 protocol (10x Genomics). Library quality was evaluated using a 2100 Bioanalyzer (Agilent) before sequencing on an Illumina NovaSeq 6000, targeting a sequencing depth of approximately 30,000 read pairs per nucleus.

### Joint snRNA-seq and snATAC-seq (snMultiome) sample and library preparation

Frozen samples were thawed at 37°C and resuspended in NP40 Lysis Buffer. Tissue dissociation was performed using a 70 µm MACS SmartStrainer (Miltenyi Biotec) with five strokes using a loose pestle followed by ten strokes with a tight pestle. The nuclei suspension was sequentially filtered through 70 µm and 40 µm Flowmi cell strainers (Bel-Art), stained with 7-AAD (1:1000 dilution, Invitrogen 15239004), and purified via fluorescence-activated nucleus sorting, with at least 200,000 intact nuclei collected per sample. Nuclei were permeabilized with 0.1X Lysis Buffer and subsequently resuspended in Diluted Nuclei Buffer as per the 10x Genomics guidelines, achieving a final concentration of approximately 5,000 nuclei/µl. All buffers used were freshly prepared. Libraries were assessed for quality using a 2100 Bioanalyzer (Agilent) and sequenced on a NovaSeq 6000 (Illumina), targeting a sequencing depth of approximately 50,000 reads per nucleus for RNA and 30,000 read pairs per nucleus for ATAC sequencing.

### Whole exome sequencing

Whole-exome sequencing libraries from the primary tumor of the patient with DMG and abdominal metastasis were generated using the KaPa hyperprep kit in combination with the HyperExome capture kit (Roche). Libraries were sequenced on a NovaSeq (Illumina) using 2x150 bp paired-end sequencing. Reads were mapped against the hg38 human genome using BWA with default parameters. Obtained CRAM files were converted into BAM files using samtools^1^, and genome-wide copy number calling was performed using CNVkit^2^ with default parameters.

### Cyclic immunofluorescence

Cyclic immunofluorescence was carried out on cryopreserved brain sections. Sections were thawed at room temperature for 30 min, post-fixed in 4% paraformaldehyde for 10 min, and rehydrated with three PBS washes. Quenching was performed in PBS containing 10 mM glycine for 20 min, followed by additional PBS washes. Samples were permeabilized with PBS/0.2% Triton X-100 for 10 min and encircled with a hydrophobic barrier. After three short PBS washes, sections were blocked in MACSima™ Running Buffer (Miltenyi Biotec) supplemented with 10% donkey serum (Merck) for 1 h at room temperature. Primary antibodies were diluted in MACSima™ Running Buffer and incubated for 2.5 h at room temperature in a humidified chamber with gentle agitation. After three PBS washes, secondary antibody (donkey anti-rabbit AF647, Thermo Fisher, 1:200) was applied for 1 h in the dark. Sections were rewashed (3 × 5 min in PBS) and mounted with SlowFade™ Gold antifade medium (Invitrogen). Imaging was performed on a Leica DMi8 Thunder microscope using a 20×/0.8 NA HC PL APO objective. For subsequent staining cycles, coverslips were gently removed in a PBS bath, and sections were washed three times (5 min each). Bound antibodies were stripped using elution buffer (Lunaphore, BU07) for 3 min, followed by another series of PBS washes (3 × 5 min). The next cycle began with blocking. Antibodies used: Round 1 - CFAP54 (Bio-Techne, NBP1-90737, 1:100); Round 2 - H3K27M (Abcam, ab190631, 1:200).

## QUANTIFICATION AND STATISTICAL ANALYSIS

### Public data collection

#### Data included in the pHGGmap (discovery cohort)

Raw data (FASTQ files) from sc/nRNA-seq, snATAC-seq, and snMultiome samples from the Jessa *et al.*^3^ and snATAC-seq samples from Liu *et al*.^4^ were downloaded from the European Genome-phenome Archive (EGA) (accession numbers: EGAS00001005773 and EGAS00001007035, respectively) with previous approval from the Data Access Committee (DAC).

#### Reanalyzed tumor data used for validations (validation cohort)

TPM normalized scRNA-seq data from Filbin *et al*.^5^ was accessed at https://singlecell.broadinstitute.org/single_cell/study/SCP147/single-cell-analysis-in-pediatric-midline-gliomas-with-histone-h3k27m-mutation and only provides filtered cells. sc/nRNA-seq from Liu *et al*.^4^ was downloaded from the Gene Expression Omnibus (GEO) database as TPM normalized count matrices under accession number GSE184357 and provides matrices for the filtered cells used in their study (with available annotation) and the unfiltered ones. scRNA-seq from LaBelle *et al*.^6^ was downloaded from the GEO database as TPM-normalized count matrices under accession number GSE227983 and provides matrices only for the filtered cells used in their study (with available annotations). Raw gene x cell matrices from scRNA-seq from DeSisto *et al*.^7^ were accessed in GEO under the accession number GSE231860. Seurat objects containing raw gene x cell matrices from snRNA-seq from Sussman *et al.*^8^ were downloaded at the Human Tumor Atlas Network (HTAN) data portal: https://data.humantumoratlas.org/.

For all tumor public datasets, donor inclusion criteria included pediatric patients according to the definition of the American Academy of Pediatrics (0-21 years old)^9^ and only tumors defined on the recent WHO2021 for brain tumors in the category of Pediatric-type diffuse high-grade gliomas defined by H3 status including only cases that were Diffuse midline glioma, H3 K27-altered and Diffuse pediatric-type high-grade glioma, H3-wild-type and IDH-wild-type.

#### scRNA-seq human developmental atlases

Processed data from Braun *et al*.^10^ (includes raw gene x cell matrices) was accessed in the provided link in their GitHub repository https://github.com/linnarsson-lab/developing-human-brain. The annotation in the original paper was too coarse to compare the granular cell states during early brain development and cancer MPs. The study by Mossi Albiach *et al*.^11^ generated higher-resolution annotations, particularly of the radial glia and glioblast populations, further identifying pure glioblasts, OPCs, preOPCs, pre-astrocytes, and pure RG. This annotation was kindly provided by Alejandro Mossi Albiach and incorporated into the original dataset. Processed data from Wang *et al*.^12^ (includes raw gene x cell matrices) was accessed in the NeMO archive at https://assets.nemoarchive.org/dat-oiif74w. Metadata was downloaded at https://cell.ucsf.edu/snMultiome. Processed data of the Li *et al*.^10^ was accessed at https://ngdc.cncb.ac.cn/bioproject/browse/PRJCA003628. Metadata detailing all different subpopulations displayed in the publication was kindly uploaded to the same repository by the corresponding author upon contact (OMIX005402, OMIX005437). For the *in silico* mix of the developmental brain dataset from the Linnarson lab with a detailed atlas of the adult brain published by Siletti *et al*.^13^, we downloaded the processed adult brain dataset (including raw gene x cell matrices) from the links provided in their GitHub repository https://github.com/linnarsson-lab/adult-human-brain.

#### snATAC-seq/snMultiome human developmental atlas

Processed data from Mannens *et al*.^14^ (includes raw peak x cell matrices) was accessed in the links provided in their GitHub repository https://github.com/linnarsson-lab/fetal_brain_multiomics.

#### Visium ST

Processed data from Ren *et al*.^15^ (includes raw peak x cell matrices and spatial imaging/coordinates) was downloaded in the GEO under accession number GSE194329, and only DMG samples were included in our analysis. Data from Collot *et al*.^16^ was shared before publication, and access to the data will be provided in that study. Processed files generated in the Kordowski *et al*. study^17^ were available through the GEO database under accession number GSE280990.

#### CosMx ST

Raw data from the van den Broek *et al*.^18^ was provided directly from the author before publication. Upon acceptance, data will be available to download at https://zenodo.org/records/1422594. Only samples from DMG donors were used for analysis.

### sc/nRNA-seq and snMultiome (GEX) data processing and quality control

For the pHGGmap, FASTQ files from our cohort and Jessa *et al*.^3^ were mapped using CellRanger version 6.1.2 (GRCh38; Ensembl98). We used the argument ‘--include-introns’ for all samples (either cells or nuclei). The samples generated using the Multiome kit were mapped using CellRanger-ARC version 2.0.2.

To correct ambient RNA, we used CellBender^19^. We decided to use this tool since it showed better ambient RNA correction in neuronal tissues^20^. For the multiome samples, we used the output from CellRanger-ARC (h5 contains both gene expression and peak matrices) and retained only the gene expression data as input for CellBender. A few challenging samples from Jessa2022, where the barcode rank plot did not show a clear curve drop, required lowering half the value of the learning rate (default: 1e-4) and increasing the number of droplets (40,000 - default:30,000) to reach convergence in the learning curve. We kept the cell barcodes inferred by Cellbender. No cells were filtered in this step.

We estimated the presence of doublets using two tools to determine doublets accurately: DoubletFinder^21^ and scDblFinder^22^. Since we had not yet filtered the cells, we divided the data into two main groups (down and high quality), initially based only on having fewer or more than 200 genes per cell, respectively. After running the doublet detection tools, we added the estimates in the metadata, and those that were not included in this analysis were labeled as ‘Not assessed’. We combined the score of both methods, as explained in https://bioconductor.org/packages/devel/bioc/vignettes/scDblFinder/inst/doc/scDblFinder.ht <u>ml</u>, calculating both the mean of the scores and the Fisher *p*-value aggregation of 1-score. To determine the threshold, we used cells assigned as singlets and doublets by both tools (‘ground truth’). Then, we calculated the receiver operating characteristic (ROC) curve and obtained the threshold that maximizes the Youden’s index on the ROC curve, using the ‘coords()’ function from the pROC package.

After filtering our cell doublets, we used MAD (median absolute deviation) thresholding to identify low-quality cells and mark them as outliers. We calculated two groups of outliers for the metrics ‘log1p_total_counts’, ‘log1p_n_genes_by_counts’, and ‘pct_counts_in_top_20_genes’ using MAD thresholds of 5 and 3. For the metrics ‘pct_counts_mt’ and ‘pct_counts_ribo,’ we used 3MAD. In addition, we excluded cells with a percentage of MT genes higher than 30%. We retained only protein-coding genes for downstream analysis. We generated four groups based on the MAD threshold and a minimum number of genes (100 or 200). We assessed among the different groups whether the different thresholds preserve or exclude cells that express classical markers of cells in the tumor (particularly immune cells, which feature a lower number of genes). We did not find a dominant population with lower cell transcripts but rather a mix of different transcriptomic signatures of different cell types (malignant, immune, vascular, etc.), so we decided to include cells that express at least 200 genes and within 3MAD.

### snATAC-seq and snMulltiome (ATAC) data processing and quality control

FASTQ files were mapped using CellRanger-ATAC version 2.1.0 (GRCh38; Ensembl98). The samples generated using the Multiome kit were mapped using CellRanger-ARC version 2.0.2. Data processing and analysis were performed using the Signac^23^ and Seurat^24^ packages, leveraging parallelization to ensure computational efficiency. When merging multiple single-cell chromatin datasets, peak calling may have been performed independently in each dataset, resulting in non-identical sets of genomic intervals. To address this, we first unified these datasets into a common peak set, applying functions from the GenomicRanges package to merge intersecting peaks, and thereby creating a single, standardized reference, as explained in https://stuartlab.org/signac/articles/merging. This approach provides a shared genomic coordinate framework against which all datasets can be quantified, ensuring that each cell is compared across the same features. Initial filtering was performed to include only cells with at least 500 fragments. After filtering and quantifying the unified peak set, each dataset was converted into a ChromatinAssay object. Finally, these standardized objects, each retaining unique sample identifiers, were merged into a single integrated Seurat object to facilitate downstream comparative analyses.

We also employed a flexible, sample-specific filtering strategy, rather than a universal cutoff, to remove low-quality cells and ensure that each dataset retained only high-quality cells suited to its inherent variability. Specifically, we examined several key metrics, including the total number of unique fragments overlapping called peaks (peak_region_fragments), the percentage of reads mapping to peak regions (pct_reads_in_peaks), the ratio of fragments overlapping blacklisted genomic regions (blacklist_ratio), nucleosome signal as a proxy for chromatin accessibility complexity (nucleosome_signal), and the transcription start site (TSS) enrichment score, which reflects signal-to-noise ratio in accessible promoter regions. For each dataset, thresholds for these metrics were determined empirically, typically requiring more than 1000-3000 unique fragments per cell, more than 5-10% reads in peaks, a blacklist ratio below 0.05-0.2, a nucleosome signal below 1.5-3.0, and a TSS enrichment score above 2.0. Median values for these metrics within each dataset generally fell well above the chosen cutoffs, ensuring that only cells demonstrating robust chromatin accessibility signals and minimal technical artifacts were included in downstream analyses.

After filtering, we employed two orthogonal computational strategies to identify and score potential doublets within the single-cell ATAC-seq and multiome datasets. First, we used the scDblFinder package^22^. Additionally, we applied a coverage-based doublet-detection method derived from the AMULET algorithm^25^, re-implemented in scDblFinder, which leverages fragment coverage properties to identify aberrant profiles potentially arising from cell multiplets. To consolidate these two independent scoring methods, we combined their *p*-values using Fisher’s method, thus generating a consensus doublet probability for each cell. A > 0.05 threshold of the combined metric was then chosen to exclude putative doublets while retaining the maximum number of high-quality, biologically meaningful singlets.

In addition to utilizing Signac and Seurat for initial data processing and quality control, we incorporated ArchR^26^ to enhance our analysis of single-cell ATAC-seq and multiome datasets. ArchR was employed to perform genome-wide tiling of accessible chromatin regions using 500-bp bins. ArchR’s tiling strategy complemented the peak calling performed by Signac by providing an alternative resolution for identifying accessible regions, which was particularly useful for capturing subtle chromatin accessibility variations that single-peak approaches might miss. To initiate this process, Arrow files were generated by inputting the fragment files obtained from CellRanger outputs into ArchR’s ‘createArrowFiles()’ function. This step included the creation of a TileMatrix, which records insertion counts across the predefined genomic bins. Quality control within ArchR was synchronized with the filtering criteria established in Signac/Seurat to maintain consistency across analysis platforms. Specifically, only cells that had passed the initial filtering steps based on fragment counts, peak region percentages, blacklist ratios, nucleosome signals, and TSS enrichment scores were retained for downstream ArchR analysis. This approach ensured the ArchR object reflected the high-quality, biologically meaningful singlets identified earlier. The integration of ArchR with Signac/Seurat allowed for the seamless transfer of cell identifiers, enabling comprehensive comparative analyses across different methodological pipelines.

### Mosaic data integration and benchmarking

Integrating single-cell datasets produced by multiple omics technologies is essential for accurately characterizing cellular heterogeneity, yet poses substantial difficulties when some data sources measure only one modality while others measure two or more. This “mosaic integration” scenario complicates modality alignment and batch effect removal, as the features and their coverage can vary significantly among datasets. To address these challenges, we explored several frameworks for integrating partially paired multi-omic data. For testing and computational efficiency, we used a subset of the dataset, down-sampling each modality to approximately 20,000 cells. First, we applied MultiVI^27^ by unifying single-modality (RNA-only or ATAC-only) and multi-omic (RNA+ATAC) datasets. We ensured consistent feature ordering (genes before peaks) and specified the primary batch annotation as “SampleID” or “Modality” for improved correction of batch-specific variation. The MultiVI model then learned a shared latent representation that bridges gene expression and chromatin accessibility profiles. Next, we tested GLUE^28^, which constructs a regulatory prior linking genomic regions to their target genes; we performed a joint variational inference approach that leverages these links for more biologically guided alignment of scRNA-seq and snATAC-seq embeddings. We also employed Multigrate^29^, wherein we organized separate RNA and ATAC layers and used different loss functions to align the modalities in a common low-dimensional space. For GLUE and Multigrate, we use “Modality” as the batch key for correction and, alternatively, add the “Batch” as a covariate. Finally, to systematically benchmark the performance of these methods, we relied on the scIB pipeline^30^ (metrics measured on the GEX). This pipeline quantifies the extent of batch correction (measured by metrics such as kBET or ASW) alongside the preservation of biological signals (e.g., cell-type separability and trajectory conservation). Through these multi-criterion evaluations, we found that all mosaic integration strategies effectively mitigated batch effects to varying degrees, with GLUE offering the best overall balance between robust correction and retention of biologically meaningful structure in our multi-omic single-cell dataset.

In the final GLUE integration (discovery cohort), we first identified a set of 2,000 highly variable genes (HVGs) in the RNA data (using ‘sc.pp.highly_variable_genes’ and a batch correction key) and retained roughly 40,000 highly variable peaks in the ATAC data (filtered by ‘min_mean=0.02’ and ‘min_disp=0.5’). We then performed Latent Semantic Indexing (LSI) on the ATAC profiles, reducing each sample to 101 dimensions and discarding the first dimension (often correlated with sequencing depth). For the RNA data, we carried out normalization, log transformation, and PCA on the 2,000 HVGs. Next, we supplied the gene coordinates (obtained via ‘scglue.data.get_gene_annotation’) along with the peak coordinates to ‘scglue.genomics.rna_anchored_guidance_graph’ to construct a regulatory prior linking putative peak-gene pairs. With the RNA and ATAC data thus annotated and embedded, we trained a ‘PairedSCGLUEModel’ by calling ‘scglue.models.fit_SCGLUÈ, specifying our annotated data and the regulatory prior graph. This model learned a shared latent space (X_glue) that integrates expression and accessibility while leveraging known regulatory relationships. Finally, we performed neighbor graph construction and Uniform Manifold Approximation and Projection (UMAP) dimensionality reduction on the integrative embedding.

### Detection of malignant cells through inference of CNV in the sc/nRNA-seq and snMultiome (GEX)

To distinguish normal from malignant cells, we inferred copy number variations (iCNVs) using a haplotype-aware approach from the Numbat package^31^, which integrates gene expression, allelic ratios, and population-derived haplotype information to detect allele-specific CNVs. Before proceeding with the iCNV analysis, we generated a preliminary cell-type annotation by mapping our dataset to the publicly available GBmap reference^32^ (via the Azimuth workflow^33^). Briefly, we performed standard preprocessing (normalization, log transformation, principal component analysis) and identified anchor features between our query data and the reference. We then transferred the reference-based labels to each query cell, obtaining predicted annotations at multiple levels of granularity. Using these transferred annotations, we identified normal-like cell populations (e.g., immune, vascular, and mature oligodendrocytes) to serve as an internal reference for copy number calling. We generated phased allele counts for each sample and then ran Numbat with default parameters. Cells displaying significant aneuploidy (i.e., subclonal copy number events) were classified as malignant, while those with diploid-like profiles were classified as normal. For a subset of challenging samples where the default settings yielded inconclusive CNV calls, we modified parameters such as ‘init_k’, ‘max_entropy’, or ‘min_LLR’ to enhance detection resolution.

### Evaluation of variance explained by biological and technical factors

We assessed the proportion of transcriptional variance attributable to both biological and technical covariates using a linear mixed-model framework adapted from the approach described in Madissoon *et al*.^34^. Briefly, we first subset cells (e.g., 30% of the total dataset) to manage computational load, removing genes with excessive sparsity (i.e., those expressed in fewer than 5% of cells). We then log-transformed total UMI and gene counts in each cell to include them as covariates alongside other sample-level annotations (e.g., batch identifiers, location, and tumor subtype) in a linear mixed model. This model decomposes the observed gene expression variability into contributions from each factor, yielding an “explained variance” estimate per factor. By comparing these fractions across the entire dataset or stratified subsets (e.g., nuclei versus whole-cell profiles), we quantified the relative influence of technical parameters (e.g., total UMI count) versus biological factors (e.g., tumor subtype, sample identity) on gene expression variation (see **Technical note**).

### Differential Gene Expression Analysis

Differential gene expression (DGE) analyses were conducted using the Seurat package, applying the Wilcoxon rank-sum test via the ‘FindMarkers()’ function. Unless otherwise specified, comparisons were performed between defined identities (e.g., treatment groups, cell states, tissue origins), with thresholds set at log₂ fold-change ≥ 1 and adjusted p-value < 0.05 (Benjamini-Hochberg correction). For selected comparisons (e.g., single-cell vs. single-nucleus), we extracted the top upregulated genes in each group and summarized their expression using ‘AggregateExpression()’ on log-normalized data. Global expression concordance between groups was further evaluated by calculating Pearson’s correlation coefficients and fitting a linear regression model, with the coefficient of determination (*R*²) reported. All visualizations and correlation metrics regarding the differential expression analysis of single cells vs. single nuclei are provided in the corresponding supplementary materials (see **Technical note**).

### Defining meta-programs (MPs) through NMF-based within-tumor variability

To systematically capture intratumoral heterogeneity within each cell compartment (malignant & myeloid), we followed a methodology adapted from Gavish *et al*.^35^, which employs a non-negative matrix factorization (NMF) within each tumor separately. This approach reduces sensitivity to batch effects and enhances the detection of gene correlations within programs across different donors, studies, and techniques, allowing the definition of biologically meaningful gene programs. First, we subset each cell compartment in each sample, filtered out known mitochondrial and ribosomal genes, and applied NMF over a range of ranks (*k*=2-10). For each NMF factor, we extracted the top 50 genes by loading weight, thereby generating a set of “NMF programs.” We then designated an “NMF program” as robust if it consistently emerged for multiple values within a given tumor, was non-redundant within that tumor (based on gene overlap), and shared at least 20% of its top 50 genes with a program from another tumor. After consolidating these robust programs across all tumors, we computed pairwise overlap (Jaccard-like similarity based on gene intersection) between programs and hierarchically clustered them. Clusters that surpassed minimal thresholds (e.g., at least 10 overlapping genes, a minimum cluster size of five programs) were considered robust meta-programs (MPs). Each MP was defined by a consensus set of 50 genes most frequently shared among the programs in that cluster or, when needed, supplemented with high-scoring genes from the member programs. This final set of MPs provided a distilled catalog of recurring transcriptional modules driving intratumoral variability across multiple tumors.

### Meta-program annotation and cross-referencing

Initially, for each MP, we first computed a per-cell score using the UCell^36^ module-scoring approach, measuring how strongly each cell expresses that MP. We then mean-centered these scores (subtracting the global average per MP) and assigned each malignant cell to the MP with the highest normalized score. Cells whose top score was not sufficiently dominant (i.e., the second-highest MP score exceeded 85% of the highest) were labeled as “unresolved,” indicating that those cells did not show clear enrichment for a single MP. Next, to characterize and annotate the MPs obtained from the NMF-based approach, we employed several complementary strategies:

1. Correlation with Published Gene Signatures: To interpret the biological relevance of each MP, we followed an approach similar to the one implemented by Qin *et al*.^37^. We compared the MP gene composition and module scores against curated or published gene signatures relevant to DMGs, other brain tumors, and pan-cancer contexts (malignant signatures^3,4,35,38–43^ and myeloid signatures^35,44–48^). We calculated UCell scores for these published signatures. Then, we performed Pearson correlation on the (scaled) module scores to build correlation matrices between MPs and known cancer-related or lineage-specific gene sets. Hierarchical clustering on these correlation matrices helped us group MPs by functional relatedness and detect parallels with existing oncogenic or lineage programs.
2. Reference-Mapping to External Single-Cell Atlases: Finally, we integrated domain knowledge from cell-type-specific references to further annotate the MPs and/or cell (sub) types. Specifically, we built refined human single-cell references from publicly available datasets, retaining relevant cell types and subtypes (to define malignant MPs^10,12^ and myeloid MPs^46,47,49,50^). Then, we applied the Azimuth pipeline to compare malignant cells against these reference atlases. The mapping results provided additional insight into the similarities in cell lineages for e MP.

### Gene Set Enrichment and Gene Ontology Analysis

We performed gene ontology (GO) enrichment analysis and hallmark gene set enrichment using both classical over-representation and curated molecular signature databases. Marker genes for each malignant program were obtained using Seurat’s ‘FindAllMarkers’ function, with positive markers filtered by adjusted *p*-value and ranked by log₂ fold change. For GO Biological Process (BP) enrichment, we used the clusterProfiler package, together with org.Hs.eg.db for gene annotation. Top 50 upregulated genes per program were analyzed using ‘groupGO()’ and ‘enrichGO()’ functions, specifying “BP” as the ontology. Redundant GO terms were summarized using semantic similarity-based clustering via ‘pairwise_termsim()’ and visualized with ‘treeplot()’. For hallmark pathway analysis, we used the enrichR package, querying the MSigDB Hallmark 2020 collection against the top program-specific genes. Enriched terms were visualized using the ‘do_TermEnrichmentPlot()’ function from the SCpubr package^51^.

### Diffusion Map and Gene Expression Dynamics

To display the developmental-like hierarchy and visualize the continuum of cancer states, we applied diffusion map analysis on GLUE-integrated embeddings. For the large pHGGmap dataset (>250,000 cells), we used Scanpy’s ‘sc.tl.diffmap’ function with default parameters to compute diffusion components. To depict cancer state density across the diffusion space, we employed ‘sc.pl.embedding_density’, allowing visualization of the spatial occupancy of each transcriptional program within the manifold. For smaller datasets (<50,000 cells), we applied diffusion map dimensionality reduction using the destiny R package^52^ and computed via ‘DiffusionMap()’. The resulting eigenvectors were extracted and stored as a custom dimensional reduction. To assess dynamic gene expression patterns along diffusion trajectories, we ordered cells by their diffusion component values and plotted the normalized expression of selected genes. Expression values were z-score normalized across cells, and heatmaps were generated using the ComplexHeatmap package^53^.

### Bulk-pseudobulk comparison and custom peak enrichment

To further elucidate the chromatin accessibility states of malignant cells, we used ArchR to (1) generate pseudo-bulk profiles within our snATAC-seq data, (2) call reproducible peaks, and (3) perform overlapping enrichment analyses with bulk and custom peak sets (including fetal neuroglial pseudo-bulks derived from the Mannens *et al*.^14^ dataset):

1. Pseudo-Bulk Generation: We grouped cells by malignant program (MP) labels and then applied ArchR’s ‘addGroupCoverages()’ function. When a cell grouping had limited representation, ArchR sampled cells with partial replacement based on the provided sampling ratio to produce the desired pseudo-bulk coverage.
2. Peak Calling and Marker Peaks: Using the pseudo-bulk replicates, we ran ‘addReproduciblePeakSet()’—using MACS2—to identify reproducible peaks. We next computed a ‘PeakMatrix’. We subsequently identified cluster-specific marker peaks via ‘getMarkerFeatures()’, selecting significant peaks enriched across all different MPs, after accounting for biases in total fragment counts and transcription start site enrichment.
3. Motif and Bulk ATAC Annotations: To interpret these marker peaks, we annotated them with known motif databases ‘addMotifAnnotations(motifSet = “cisbp”)’ and tested for overlaps with a curated library of bulk ATAC-seq peak sets ‘collection = “ATAC”}’. This allowed us to assess the presence of canonical regulatory motifs and compare our single-cell-resolved peaks to established bulk accessibility datasets.
4. Custom Enrichments Using Fetal Neuroglial Reference Peaks: Beyond the curated annotation sets, we performed custom enrichment analyses to explore lineage or developmental relevance. Specifically, we incorporated fetal neuroglial pseudo-bulk peaks from the Mannens *et al*. dataset^14^, which were derived by merging cellranger output fragment files from multiple fetal samples into a single file, using a custom Python pipeline to rename barcodes (prefixing them by sample) and filter only those matching the annotated reference. Then, we split the merged file by cell type via ‘snATAC_fragment_tools’ (https://github.com/aertslab/snATAC_fragment_tools?tab=readme-ov-file), retaining only neuroglial lineages (e.g., radial glia, neural progenitors, oligodendrocyte precursors, and astrocyte-like cells). Lastly, we generated a final set of bed-like fragment files for each cell type. These fetal neuroglial fragment files were loaded into ArchR ‘addPeakAnnotations()’ as a custom annotation “fetal,” enabling us to run the ‘peakAnnoEnrichment()’ function for each group of marker peaks. The resultant enrichment heatmaps highlighted which malignant programs share accessibility patterns with specific fetal glial subpopulations.

### SCENIC+ analysis

SCENIC+^54^ was used to infer gene regulatory networks (GRNs) from our integrated sc/nRNA-seq and snATAC-seq datasets. We first pre-processed sc/nRNA-seq data in Scanpy, normalizing read counts and transforming them with a log scale. For the snATAC-seq data, we employed pycisTopic^54^ to create a cisTopic object, model latent topics, and impute accessibility scores. Importantly, we included all cell type profiles (i.e., malignant and non-malignant cells) so that SCENIC+ had sufficient diversity to define robust transcription factor-driven enhancer modules. We then ran the SCENIC+ Snakemake pipeline, providing per-cell gene expression and the cisTopic-based accessibility matrices, along with sets of candidate genomic regions (e.g., marker peaks, topic-specific peaks). SCENIC+ generated “direct” and “extended” eRegulons, linking candidate transcription factors to putative target genes via co-accessible regulatory elements. We computed eRegulon AUC scores (reflecting per-cell regulon activity) and derived “regulon specificity scores” to highlight eRegulons enriched in the malignant states. Finally, we visualized the resulting eRegulons and GRN networks in Python (with NetworkX) and exported them to Cytoscape for further graphical annotation.

### Epigenetic plasticity analysis

Following the strategy described by Burdziak *et al*.^55^, we quantified epigenetic plasticity by evaluating how snATAC-seq profiles can simultaneously support multiple, distinctly defined transcriptomic cell states. Specifically, we:

1. ArchR alternative processing and feature selection: Initially, we reprocessed snATAC-seq data through a modified ArchR pipeline^56^ that targets nucleosome-free (NFR) fragments (fragment length < 147) instead of utilizing all fragments as the default does. This approach is also more compatible with the metacell algorithm, increasing the detection of regulatory elements.
2. Metacell construction and aggregation (SEACells): Given the inherent sparsity of snATAC-seq signals, we aggregated similarly profiled cells into “metacells” to stabilize estimates of cell-specific chromatin landscapes. Using the SEAcells framework^56^, we subdivided the dataset into archetypal “metacells,” each representing a distinct local region of the phenotypic manifold (as defined by the shared nearest-neighbor graph in the GLUE space). This produces aggregated (meta) profiles with improved signal-to-noise ratios for epigenomic features while still maintaining relatively fine-grained resolution of subpopulations.
3. Corresponding sc/nRNA-seq metacells: In parallel, we also sampled sc/nRNA-seq data to produce a comparable number of metacells, employing SEAcells on the transcriptomic space (GLUE). By constructing matching epigenomic and transcriptomic metacells, we aligned heterogeneous cell states from both omics layers.
4. Logistic regression classifier for cross-modal predictions: To estimate whether each epigenomic metacell could “support” multiple distinct transcriptomic identities, we trained a multi-class logistic regression model on the transcriptomic metacells as specified by Burdziak *et al*.^55^. Briefly, we treated fine-grained sc/nRNA-seq clusters as discrete classes. We then used aggregate gene expression from the RNA metacells as input features, followed by evaluating the model’s accuracy on held-out RNA metacells. Next, we substituted each epigenomic metacell’s aggregated accessibility profile (proximal to each gene) in place of the gene expression vector in the same feature space (shared gene sets, normalized appropriately). After, we obtained a probability distribution over all possible transcriptomic states for every ATAC metacell. We interpreted the entropy of this probability distribution as a “plasticity score.” If an ATAC metacell confidently mapped to a single transcriptomic class, it exhibited low plasticity. Conversely, if the model assigned non-trivial probabilities to multiple distinct transcriptomic states, it implied that the chromatin landscape primed more than one gene expression program, suggesting higher plasticity. On a per-metacell level, we computed Shannon’s entropy of the logistic regression output, capturing how “broadly” each ATAC profile spans multiple transcriptomic fates. Lastly, we compared plasticity scores across different malignant programs, highlighting which states appear most epigenetically permissive to diverse gene expression outcomes.

### CytoTRACE2-based prediction of cellular potency

To assess the developmental potential of malignant cells, we applied CytoTRACE 2 (a deep learning-based method for estimating cellular potency from scRNA-seq profiles)^57^. This tool infers both discrete potency categories (e.g., totipotent, pluripotent, multipotent, etc.) and a continuous potency score that spans 0 (highly differentiated) to 1 (developmentally “naive” or totipotent). We used the function ‘cytotrace2’ with ‘batch_size=20000’ and ‘smooth_batch_size = 3000’.

### Comparison of PM-like Marker Expression in Pediatric and Adult High-Grade Gliomas

To investigate the specificity of the PM-like malignant program to pHGG, we compared its defining gene expression signature with malignant cells from adult GB. We integrated single-cell datasets from our annotated pHGGmap and an adult GB reference (GBmap)^32^ using Seurat. Malignant cells were extracted from both datasets and merged into a single object with a unified group label (“Malignant DMG” or “Malignant GB”). The expression levels of PM-like marker genes were quantified, enabling direct comparison of expression magnitude between pediatric and adult malignant compartments.

### Cellular States Graphical Representation

To display the immunomodulatory programs identified in the myeloid compartment, we adapted the concept of “Four variable” state plots introduced by Neftel *et al*.^41^, implemented in the SCpubr package via the ‘do_CellularStatesPlot()’ function. In brief, four gene sets (x1, x2, y1, y2) are defined, and each cell’s enrichment for those sets is computed (via ‘AddModuleScorè). Each cell is then in a 2D coordinate system: along the y-axis the cell is assigned to the gene set pair (x1, x2) if its (x1 + x2) score exceeds that of (y1 + y2), or vice versa and, along the x-axis a log2 difference in scores is computed—specifically, log2(abs((x1 or y1) - (x2 or y2)) + 1), with sign determined by which gene set has the higher score. This scheme creates a “four-quadrant” style plot, where each corner highlights cells most enriched for one of the four gene sets. In our pipeline, we sometimes replaced the final plotting routine with a custom approach that uses partial code from ‘SCpubr::do_CellularStatesPlot()’ to extract per-cell coordinates. We then overlaid multi-slice “pie” representations (per cell) to depict concurrent enrichment of multiple modules.

### Label Transfer from pHGGmap to Public Datasets for Validation and Mapping Identities of Peritoneal Metastatic Cancer and Infiltrating-Myeloid Cells

To validate our findings in independent cohorts and unify annotations across different data sources, we re-annotated multiple publicly available sc/nRNA-seq datasets using a reference-based mapping approach. In particular, we began by loading our pHGGmap reference (discovery cohort) containing the curated annotation. For each public dataset (Filbin *et al*.^5^, Liu *et al*.^4^, DeSisto *et al*.^7^, Labelle *et al*.^6^, and Sussman *et al*.^8^), we performed standard preprocessing steps within Seurat (normalization, identification of variable features, scaling, PCA-based dimensionality reduction) to obtain a query object. We applied Seurat’s ‘FindTransferAnchors()’ function, specifying the pHGGmap as the reference and our query dataset as the target. Using the resulting anchor set, we invoked ‘TransferData()’ to predict cell-level annotations matching those in the reference. We further integrated embeddings via ‘IntegrateEmbeddings()’ and projected each query dataset onto the reference UMAP space through ‘ProjectUMAP()’. This pipeline assigned each cell in the public dataset an annotation mirroring the pHGGmap schema, enabling direct comparisons despite differences in the sequencing platforms (for example, SMART-seq2). We repeated this procedure for all datasets, focusing only on pediatric samples (0-21 years old) with relevant diagnoses (e.g., diffuse midline glioma H3 K27-altered; Diffuse pediatric-type high-grade glioma H3-wild-type, IDH-wild-type). In certain instances (Sussman *et al*.^8^), we refined the existing authors’ annotations to align with the atlas-defined categories. We follow the same approach for the DMG case with abdominal metastasis. We mapped the cells to resolve the cancer phenotype of the metastatic cells and the immunomodulatory programs of the infiltrating myeloid cells against the programs defined in the pHGGmap.

### Poisson Linear Mixed-Effect Model for Compositional and Covariate Analyses

To quantify how molecular or clinical factors influence cell-type abundance or gene expression, we employed a Poisson-based mixed-effect model strategy, adapted from previously described frameworks^34,58^. In general, we constructed a counts-by-sample matrix from our pHGGmap metadata and tallied the number of cells of each subtype in each sample. For example, each row corresponded to a sample (e.g., individual or library), and each column to a particular cell subtype. These counts were then used as the response variable in a generalized linear Poisson mixed model. We gathered relevant clinical or technical covariates (e.g., age, treatment status, capture modality [see **Technical note**]) at the sample level. Categorical covariates were encoded as design matrices, while numeric covariates were standardized or treated as random slopes. Any repeated measurements (e.g., multiple libraries per donor) were introduced as nested or crossed random effects. We fit the model using the lme4 package’s ‘glmer()’ function (family = poisson). The model yields random-effect estimates for each covariate-celltype combination, effectively giving a log fold-change relative to some baseline. We recast these into fold changes by exponentiation. To address whether a particular factor’s effect on a cell type is robust, we computed a local true sign rate (LTSR), which gauges the probability that the effect direction (positive or negative) is correct. LTSR values range from 0 to 1, with higher values indicating greater confidence in the estimated direction. Following established thresholds, we typically labeled effects with LTSR > 0.9 as substantially affected.

### Modeling Multicellular Niches and Trajectories with the BEYOND Framework

To evaluate variation between individuals in the frequency of different cell states and analyze changes in subtype abundance dynamics and define trajectories of changes in individual cellular environments, we employed the BEYOND framework^59^. At its core, BEYOND represents each donor’s cellular landscape (summarized as subpopulation proportions) in a high-dimensional manifold, reconstructs trajectories of cellular change across participants, and then dissects both subpopulation and trait dynamics along these trajectories. First, we aggregated single-cell data at the level of biologically meaningful subpopulations (cancer and myeloid phenotypes). For each donor, we computed a subpopulation-proportion matrix in which each row corresponds to one participant and each column to the relative abundance of a single-cell-defined subpopulation. Next, we learned a manifold via PHATE^60^ to embed participants based on their cellular environments and applied graph-based community detection (Leiden algorithm) to identify coarse groupings. Using the Palantir^61^ pseudotime algorithm, we inferred trajectories on this manifold, yielding estimates of each participant’s pseudotime along one or more “branches.” For each trajectory, we fit generalized additive models to subpopulation proportions (and relevant clinical variables) as a function of pseudotime, thereby revealing how subpopulations or traits vary along each path. Finally, we clustered subpopulations into “cellular communities” by considering both their correlation across donors (co-occurrence in participant samples) and similarity in their model-fitted dynamics, effectively grouping subpopulations whose abundance changes in tandem. Donor-level proportions of each inferred transcriptional subcommunity (C1, C2) were extracted from the BEYOND-derived donor×community representation. For each donor, C1 and C2 values were normalized to the sum of C1+C2 to obtain within-module proportions. Donors were stratified by clinical status (treatment naïve vs. post-treatment). Differences in proportions between clinical groups were evaluated independently for C1 and C2 using a two-sided Mann-Whitney U test.

### Visium Spatial Transcriptomics Deconvolution

We downloaded the raw Visium outputs from the publicly available DMG cohort^15–17^. For spatial cell deconvolution, we employed the cell2location^62^ algorithm. We used pHGGmap as a single-cell reference, wherein each cell type’s average gene expression signature was estimated using a Negative Binomial regression model (‘cell2location.models.RegressionModel’). We intersected the spatial dataset’s genes with those in the reference signatures and set two hyperparameters in ‘cell2location-N_cells_per_location’ — (estimated from histology) and ‘detection_alpha = 20’ to account for technical variation in the Visium data. We then trained a model (‘cell2location.train’) for 30,000 epochs on a GPU. This generated posterior estimates of cell-type abundance for each spot (5% quantiles), which we visualized in spatial coordinates to confirm robust detection of distinct cell subpopulations. After mapping cell type abundances to each spatial spot, we defined spatial niches by performing NMF (‘run _colocation’) on the resulting cell type abundance matrix (for the Ren *et al.* and Collot *et al*. datasets). We varied the NMF factor *k* and evaluated the consistency of extracted factors (spatial co-localizations) to converge on *k*=7.

### Mapping CosMx Spatial Data with Tangram

To transfer our single-cell annotations onto the CosMx spatial transcriptomics dataset from van den Broek *et al*.^18^ and jointly explore both transcriptomic and spatial structure, we employed Tangram^63^. We removed negative-control probes and performed basic quality checks, retaining only fields of view (FOV) associated with DMG H3 K27-altered tissues. Next, we normalized and log-transformed the CosMx data (via Scanpy’s ‘sc.pp.normalize_total’ and ‘sc.pp.log1p’). We employed our pHGGmap, containing the aggregated transcriptomes used as the source of high-resolution gene expression and cell-type assignments. Using ‘tg.pp_adatas’, we identified the shared gene set between the CosMx panel and the single-cell reference. We ran ‘tg.map_cells_to_spacè in “single-cell mode,” which fits a mapping matrix, indicating the probability that each single cell occupies each spatial spot. For computational efficiency, we used a GPU environment. Tangram optimizes a cosine-similarity-based objective such that the aggregated expression of mapped single cells in each spatial voxel best matches the measured CosMx spot expression. This step provided us with a cell-by-voxel probability matrix. Tangram’s function ‘tg.project_cell_annotations’ took the single-cell-type labels and assigned them to each spatial FOV by weighting each cell’s assignment in proportion to the mapping probabilities. We thus obtained a “tangram_prediction” for each voxel, reflecting the cell type most likely to reside there. To confirm mapping quality, we examined the distribution of training gene similarity scores (cosine similarity). Although we acknowledge that the CosMx gene panel is not fully specialized for brain tumor markers, about 60% of the training genes showed high alignment (score ≥ 0.8), suggesting the gene panel and the single-cell atlas were sufficiently compatible for the cell type assignment.

### Spatial Cell Niches Detection with CellCharter

To identify cellular niches in the CosMx dataset^18^, we employed the CellCharter framework^64^. After cell mapping with Tangram, we used scVI to derive a low-dimensional latent space. Specifically, we set up the model to account for batch effects across multiple samples (‘scvi.model.SCVI.setup_anndatà with ‘batch key = samplè) and then trained scVI with default parameters. To incorporate spatial context, CellCharter uses a graph-based strategy in which each cell is a node, with edges between physically adjacent cells. We computed these adjacency relations by calling ‘squidpy.gr.spatial neighbors’ on each sample, specifying ‘coord type = generic’ and ‘spatial key = spatial_fov’. Although the default Delaunay method can sometimes link distant cells, we optionally removed spurious connections with ‘cellcharter.gr.remove_long links’. We then aggregated local neighbors’ features out to three layers of adjacency, appending each layer’s mean scVI latent vector to the cell’s 10-dimensional vector, producing a 40-dimensional. This merging of per-cell and neighborhood signals helps CellCharter detect multicellular niches more robustly. For cluster determination, we tested multiple values of *K* (the number of spatial niches) via ‘cellcharter.tl.ClusterAutok’. This procedure repeatedly clusters the data for different *Ks*, measuring solution “stability” to pinpoint stable domain counts. We selected the final niche number based on peaks in stability (*K* = 9). To interpret niche composition and relationships, we leveraged ‘cellcharter.gr.enrichment’ to analyze whether certain cell types (transferred from Tangram predictions) are enriched in each niche.

### Computing Spatial Gene Expression Signatures

We evaluated signature (module) scores for selected gene sets in the CosMx data^18^ by first mapping each cell’s expression onto a predefined gene list. Specifically, we compiled marker genes for certain states (for example, hypoxia or stress), filtered those genes to those present in the dataset, and applied ‘scanpy.tl.score_-genes’ to each sample separately. This yielded a per-cell score representing the average expression for each gene signature. Because local smoothing often improves the visualization of noisy or spotty signals in spatial data, we performed an additional neighborhood-averaging step. We called ‘squidpy.gr.spatial_neighbors’ to build a graph of local adjacency for the relevant cells, ensuring each cell was connected to a specified number of neighbors (*n* = 50). Then, we replaced each cell’s raw signature score with a weighted average of its own and its neighbors’ scores, normalizing by connectivity degrees.

### Association Between Malignant Programs and Invasivity Signature in sc/nRNAseq

To examine the relationship between malignant transcriptional programs and tumor invasiveness, we scored cells based on two gene sets: (1) the invasivity signature from Venkataramani *et al*.^65^, and (2) malignant program-specific (MP) gene modules defined via NMF decomposition in our study (e.g., RG-, OPC-like programs). Both scores were computed using Seurat’s ‘AddModuleScorè. We restricted the analysis to tumor-classified malignant cells (iCNV+). After computing scores, we visualized the association between the selected pHGG MP and invasivity score using a smoothed scatterplot (LOESS fit). Pearson correlation was computed to quantify the strength of association, with *R* and *p*-value reported. This approach was repeated for other MP modules to rank cancer states and spatial niches by their putative invasiveness potential.

### Spatial Scoring of Gene Signatures

Spatial scoring of tissue regions was performed by calculating a gene signature score per spot using a curated list of genes associated with invasiveness or metabolic-associated programs^35,65^. For each sample, spatial gene scores were computed using the ‘score_genes’ function from Scanpy. These scores were calculated individually for each tissue section to preserve within-sample variability. To ensure comparability across samples, gene scores were min-max normalized per sample based on the observed range of values. These scores were then interpreted in the context of underlying cellular composition as inferred by cell2location or other variables present in the metadata (e.g., tumor location). Specifically, for cell2location inferred cellular composition, we used the estimated cell-type abundance matrix (5% quantile of the posterior) and defined, for each spatial location, the most enriched cell type—i.e., the one with the highest predicted abundance among all modeled populations. To enhance robustness and reduce noise, only locations where the predicted abundance exceeded the 95th percentile (q95) across all spots in the sample were retained for downstream interpretation. We then aggregated the normalized gene scores by enriched cell type and sorted them by the median invasivity score.

### Identification of RG-like Subpopulations via Hypoxia and Stress Signature Scoring

To resolve transcriptional heterogeneity within the RG-like malignant program (MP4), we scored each cell based on its enrichment for two distinct gene signatures associated with tumor microenvironment adaptation—namely, a hypoxia-associated program (MP6) and a stress-related program (MP10). Signature gene sets were obtained from NMF decomposition and applied using the UCell::AddModuleScore_UCell() function with parallel computing (ncores = 64). To define discrete RG-like subpopulations, we applied a Gaussian mixture modeling approach (mclust::Mclust) to each signature score distribution. The intersection point of posterior probabilities between the two components was used as a threshold to classify cells. RG-like cells exceeding the threshold for either MP6 or MP10 were labeled as “TE-associated”, while the remainder were categorized as “Developmental-like”.

### Graph Abstraction Using PAGA

To reconstruct discrete transcriptional transitions and connectivity between malignant subpopulations from primary and metastatic (ascites) samples, and PBMCs and myeloid-infiltrating peritoneal cells, we employed Partition-based Graph Abstraction (PAGA)^66^ as implemented in Scanpy. We first computed a k-nearest neighbors graph (‘sc.pp.neighbors’) and applied ‘sc.tl.pagà using clusters as the grouping variable. The resulting abstracted graph was visualized using ‘sc.pl.pagà, with nodes representing clusters and edge weights encoding inter-cluster connectivity based on transcriptomic similarities.

### Trajectory Inference Using Palantir

To investigate dynamic transcriptional changes, we performed pseudotime and lineage inference using the Palantir framework^61^. First, we computed a diffusion map representation using ‘palantir.utils.run_diffusion_maps‘, followed by the construction of the multiscale manifold with ‘palantir.utils.determine_multiscale_spacè. Gene expression values were denoised using MAGIC imputation^67^ (‘palantir.utils.run_magic_imputation’). The start cell for trajectory computation was selected from the primary tumor edge cluster, and terminal states were inferred from representative cells within distinct subclones, based on clustering and spatial UMAP positions. Trajectory computation was performed using ‘palantir.core.run_palantir’ with 500 waypoints, capturing cellular transitions toward distinct terminal phenotypes. Lineage branches were defined using ‘palantir.presults.select_branch_cells’. Entropy-based pseudotime progression was displayed alongside branch-specific fate probabilities. To characterize transcriptional dynamics along inferred trajectories, we used ‘palantir.presults.compute_gene_trends’, leveraging MAGIC-imputed values. Gene trend plots and heatmaps were generated for representative markers of distinct phenotypic states. Visualizations were generated using ‘palantir.plot.plot_gene_trend_heatmaps’.

### Differentially Accessible Regions and Transcription Factor Motif Analysis

The processed ArchR project containing primary and metastatic malignant cells was used to identify Differentially Accessible Regions (DARs) with ‘getMarkerFeatures()’ using the PeakMatrix and Wilcoxon rank-sum test (‘testMethod = “wilcoxon”’), while controlling for technical biases (‘bias = c(”TSSEnrichment”, “log10(nFrags)”)’). We focused on comparisons between primary AC-like, primary OPC-like, and ascites RG-like populations. A heatmap of DARs was generated using ‘markerHeatmap()’ with FDR ≤ 0.1 and log2 fold change ≥ 0.5 to visualize global accessibility differences. To identify candidate regulators, motif enrichment analysis was performed by annotating peaks with ‘addMotifAnnotations()’ (cisBP database) and using ‘peakAnnoEnrichment()’ on DARs specific to the RG-like metastatic population. Enrichment scores were ranked and visualized using ggplot2. To validate the functional engagement of these motifs, we performed footprinting analysis using ‘getFootprints()’ and visualized normalized footprints with ‘plotFootprints()’ (bias-corrected using the Subtract method).

### Program overlap and similarity analysis

To quantify the extent of shared genes between malignant and myeloid meta-programs, we computed pairwise gene-set overlaps between all program pairs. For each malignant program A and myeloid program B, we calculated the intersection size, the union size, and the Jaccard similarity index. To assess whether overlaps exceeded what would be expected by chance, statistical significance was evaluated using a one-sided hypergeometric test, with the universe defined as the union of all unique genes appearing in any of the compared programs. *p*-values were corrected for multiple testing using the Benjamini-Hochberg procedure, and adjusted values (FDR) are reported. Only program pairs with FDR < 0.05 were considered significantly overlapping.

## Technical note - Single-Cell vs. Single-Nucleus Profiling

We evaluated how isolation protocols (i.e., fresh cells vs. frozen nuclei) affect the detection of different cell types and transcriptional programs in pHGG. Single-nucleus RNA sequencing (snRNA-seq) captured cell types that are more sensitive to mechanical digestion, such as neurons and macroglia (astrocytes and oligodendrocytes), which were underrepresented in single-cell RNA sequencing (scRNA-seq) (**Figure A**). Although sample isolation and 10x Genomics kit versions introduced minor biases, these factors explained less than 1% of the overall gene expression variance (**Figure B**). Thus, combining both scRNA-seq and snRNA-seq approaches provides a more comprehensive survey of cellular diversity within the tumor.

To quantify how capture modalities could influence cell type proportions, we utilized a Poisson linear mixed model as previously described^1,2^. We observed significantly higher proportions of differentiated-like (AC) tumor cells in nuclei-derived samples. In contrast, cell-derived samples showed enriched proportions of stem-like cells (particularly RG-like) (**Figure C**). This disparity may reflect how differentiated cancer cells are more prone to damage during tissue digestion, thereby favoring their capture in nuclei-based protocols, similar to what occurs with healthy neurons and oligodendrocytes. A similar discrepancy between cell proportions in scRNA-seq and spatial transcriptomics (ST) has been reported by Liu *et al.*^3^.

We also noted differences in the detection of TAMC activation genes between scRNA-seq and snRNA-seq, consistent with previous work^4^. For example, complement- and proinflammatory-related MPs were more readily observed in cell-derived TAMCs than in nuclei-derived TAMCs (**Figures D and E**). Specific activation genes (*APOE, CST3, SPP1*) also showed higher expression levels in cell-derived TAMCs (**Figure F**). Despite these differences in expression magnitude, snRNA-seq still reliably recovered inflammatory programs, and gene expression values correlated highly (*R*² = 0.9) between cell- and nucleus-derived samples (**Figures F and G**). Collectively, our assessment suggests that while each capture modality has inherent biases, both approaches capture essential aspects of the pHGG cellular landscape.

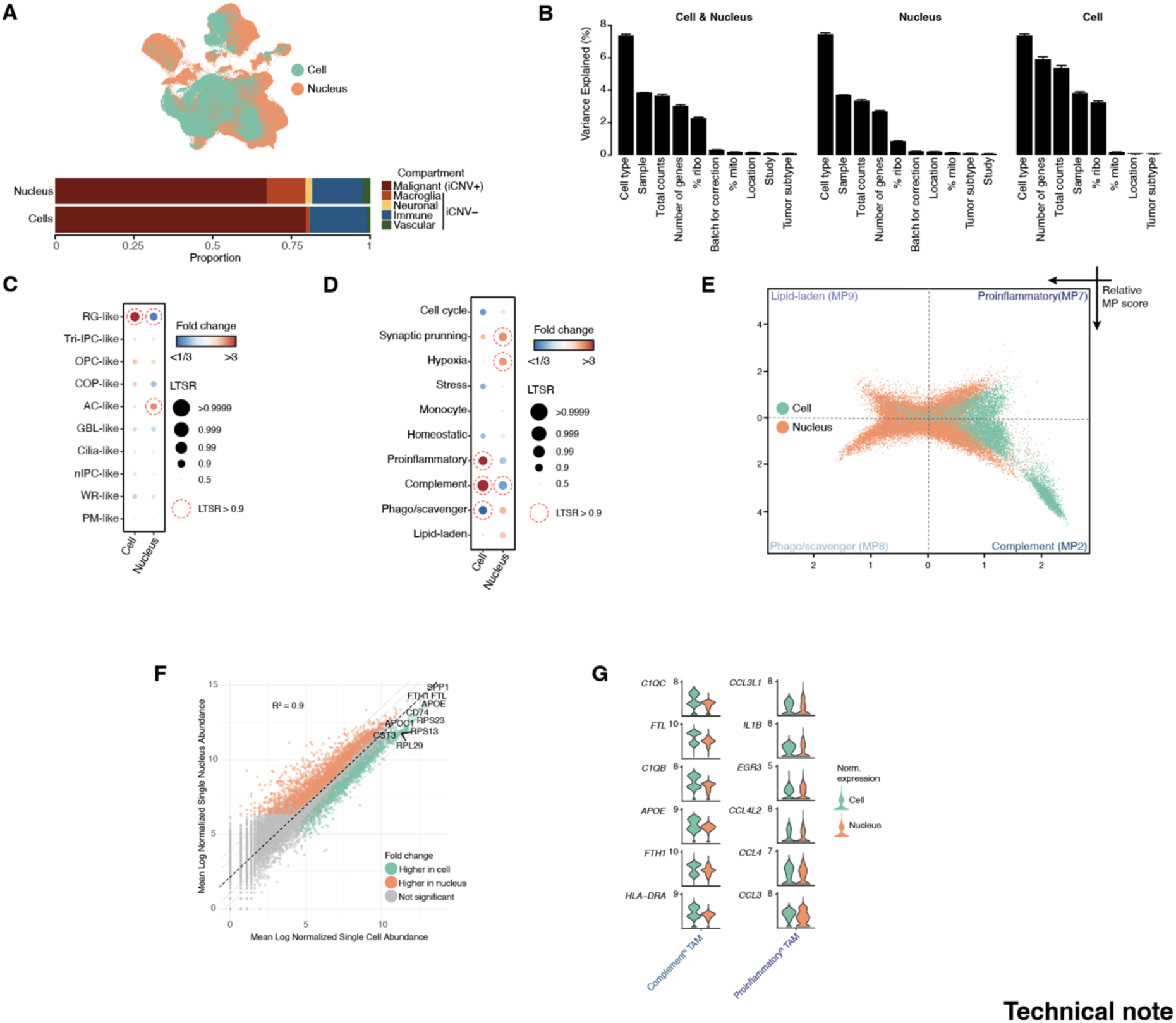
(A) Stacked barplot of the proportion of cells per each capture modality. (B) Bar charts illustrate the variance in gene expression accounted for by metadata variables across the combined sc/nRNA-seq, scRNA-seq, and snRNA-seq datasets. The error bars represent 95% confidence intervals. (C-D) Cell type proportions analysis shows log2-transformed fold change and LTSR score, depicting variations in cell cluster abundance over the capture modality. LTSR spans from 0 to 1, with a value of 1 representing a reliable estimate. (E) Representation of TAMCs’ cell states. Each quadrant represents a distinct type of immunomodulatory TAMC MP. Colored by capture modality. (F) Scatter plot of the average normalized gene expression in cells (x-axis) and nuclei (y-axis). Genes with significantly higher expression in nuclei are marked in orange (*p* adj < 0.05, fold change > 1), while genes with significantly lower expression in nuclei are indicated in green (*p* adj < 0.05, fold change < -1). *R*^2^ represents the correlation coefficient. (G) Violin plot of gene expression of selected markers for MP2 (complement) and MP7 (proinflammatory), split by capture modality.

## Notes

https://doi.org/10.5281/zenodo.17063631

